# LIF-LIFR/gp130 Survival Signaling and Connexin 47 Dysregulation upon Murine-β-Coronavirus Infection: Discovery of a Novel ERK2 Phosphorylation Site at Cx47 C-Terminal Domain

**DOI:** 10.64898/2026.09.21.753212

**Authors:** Santosh Kumar Samal, Rishika Jana, Sayar Ghosh, Anshita Mishra, Ankita Mandal, Subhajit Das Sarma, Jayasri Das Sarma

## Abstract

Oligodendrocyte (OLG) injury and loss are increasingly recognized as major determinants of demyelination and remyelination failure in multiple sclerosis (MS), alongside conventional immune-mediated processes. However, the mechanism underlying OLG injury and remyelination failure remains incompletely understood. Astrocyte-oligodendrocyte interactions mediated by Connexin (Cx)-43-Cx47 gap junction (GJ) communication and leukemia inhibitory factor (LIF)-LIFR/gp130 signaling are critical for oligodendroglial survival, homeostasis, and myelination. Despite their established role, how these pathways are altered during neuroinflammatory diseases remains unclear. To address this, we integrated a murine β-coronavirus (RSA59)-induced *in vivo* model with enriched primary oligodendrocyte precursor cell (OPC) and mature OLG cultures to investigate the effect of RSA59 infection on OLGs, Cx47-mediated GJ communication, and LIF-LIFR/gp130 Signaling. We observed that RSA59 directly targets oligodendroglial lineage cells, including OPCs and mature OLGs, and induces apoptosis in mature OLGs. Despite increased OPC abundance, RSA59 infection reduces the expression of mature and myelinating markers CNPase and MBP, raising the possibility of impaired oligodendrocyte maturation and myelination. RSA59 infection also differentially regulates Cx47-mediated GJ communication and LIF-LIFR/gp130 signaling, crucial for OLG survival and homeostasis. Furthermore, our studies demonstrate that extracellular signal-regulated kinase (ERK), a downstream effector of LIF signaling, phosphorylates the Cx47 C-terminal domain (Cx47CT) at Serine-372 residue, identifying a previously unreported mechanism. Collectively, these findings reveal that RSA59 infection disrupts interconnected oligodendroglial communication and survival pathways, providing mechanistic insights into virus-induced chronic demyelination pathology. The study further identifies Cx47-mediated GJ communication and LIF-LIFR/gp130 signaling as crucial therapeutic targets, and highlights ERK-dependent Cx47CT phosphorylation as a potential regulatory mechanism.

## Introduction

Oligodendrocytes (OLG) are highly vulnerable to immune-mediated injury, and their damage contributes to oligodendroglial loss, focal demyelination, axonal damage, and progressive neurological disability in multiple sclerosis (MS) (1–3). Although MS has traditionally been viewed as an immune-mediated disorder, accumulating evidence indicates that OLG death and damage may actively contribute to lesion formation and disease progression (4). In certain contexts, early OLG injury may precede and contribute to subsequent immune activation, indicating a bidirectional relationship between oligodendroglial damage and neuroinflammation (5, 6). Additionally, despite the recruitment of oligodendrocyte progenitor cells (OPCs) to demyelinated lesions, impaired OPC differentiation and maturation can limit effective remyelination (7). Therefore, understanding the mechanisms that regulate oligodendrocyte survival, maturation, and communication with neighboring neuroglial cells is critical for developing strategies to prevent demyelination and promote remyelination.

OLGs communicate with astrocytes through direct cell-cell contacts and via paracrine signaling involving cytokines, chemokines, extracellular vesicles, and other soluble factors (8). Among these mechanisms, connexin (Cx)-mediated gap junction intercellular communication (GJIC) enables the direct exchange of ions, metabolites, and signaling molecules (9). Astrocytes predominantly express Cx43, whereas OLGs express Cx47, which form heterotypic gap junction channels (GJCs) that facilitate small metabolite exchange, K^+^ buffering, and maintenance of myelin integrity (10). Perturbation of Cx43-Cx47 functional coupling can disrupt metabolic support and cellular homeostasis, thereby affecting OLG health and, in turn, myelin health. Loss of Cx43-Cx47 GJCs has been reported in MS lesions and is associated with distal oligodendrogliopathy and increased disease severity (11, 12). Consistently, mice lacking Cx47, Cx43/Cx30, and Cx30/Cx47 GJ proteins develop extensive white matter vacuolization and myelin abnormalities, underscoring the importance of Cx-mediated coupling for CNS homeostasis (13). Despite this, the molecular mechanisms behind Cx47 dysregulation during inflammatory demyelination remain understudied.

Cx43-Cx47 GJICs can also propagate crucial signaling molecules that regulate various oligodendroglial functions. For example, neuronal ATP stimulates astrocytic secretion of leukemia inhibitory factor (LIF), which promotes OPC maturation and subsequent myelination (14). Thus, coordinated Cx43-Cx47 communication and LIF-mediated signaling constitute an important component of astrocyte-oligodendrocyte functional coupling.

LIF is a pleotropic cytokine that belongs to the interleukin-6 (IL-6) superfamily and signals through the LIF receptor (LIFR) and gp130 receptor complex. Activation of this receptor complex stimulates several downstream intracellular pathways, including JAK-STAT, AKT, MAPK/ERK, and mTOR signaling (15, 16). LIF is generally undetectable in the healthy nervous system, but is upregulated in response to injury, suggesting an endogenous role in neuroprotection (17, 18). Studies in animal models have demonstrated that LIF-deficient mice exhibit severe myelination defects in the optic nerve and delayed OPC maturation during postnatal development (19), whereas reduced LIF receptor signaling results in accelerated experimental autoimmune encephalomyelitis (EAE) progression, increased oligodendrocyte death, and severe demyelination (20, 21). LIF can also protect oligodendrocytes from the detrimental effects of TNF-α and IFN-γ, both of which are elevated in MS (22, 23). In addition to its protective effects on OLGs, LIF exerts distinct immunomodulatory effects. In contrast to IL-6, which promotes IL-17 production by T cells and suppresses regulatory T cell (Treg) responses, LIF has been reported to enhance Treg function in MS while inhibiting IL-17 production by T cells (24, 25). Considering the relevance of LIF-LIFR/gp130 signaling in oligodendroglial biology and MS, this study further explores how viral infection modulates this protective pathway.

Studies using the murine β-Coronavirus (MHV-A59/RS59), an experimental model of virus-induced demyelination (26), have established various mechanisms altering astrocyte-oligodendrocyte communication, with a particular emphasis on astrocytic Cx43. In primary astrocytes, MHV-A59 infection leads to ER-ERGIC retention of Cx43 and downregulates its cellular expression through virus-induced competitive inhibition of microtubule-mediated intracellular trafficking (27, 28). Cx43 accumulation within the ER-ERGIC compartment induces ER stress and downregulates ERp29, a crucial chaperone for Cx43 trafficking.

Restoration of ERp29 expression, either through pharmacological treatment with 4-phenylbutyric acid (4-PBA) or exogenous overexpression, rescued Cx43 trafficking in vitro (29). Similarly, in vivo treatment with 4-PBA also stabilized ERp29 expression and thereby supported the expression of gap junction proteins Cx43 and Cx47 in the mouse CNS (30). Building on these insights from previous studies of Cx43, the present study attempts to address the existing research gaps associated with Cx47.

Previous studies have demonstrated that MHV-A59 infection induces persistent reduction of Cx47 GJs in mature OLGs within demyelinated and adjacent normal-appearing white and grey matter regions of the spinal cord (31). Similarly, reduced Cx43 expression led to the destabilisation and downregulation of Cx47 in MHVA59-infected mouse brains (28). Although the exact mechanism remains unknown, reports suggest that astrocytic Cx43 is essential for Cx47 phosphorylation and surface stabilisation in oligodendrocytes (32). Phosphorylation of Cx proteins regulates oligomerisation, trafficking, GJC formation, gating, internalisation, and degradation (33). Several studies have documented phosphorylation of the Cx43 C-terminus (Cx43CT), with recent evidence indicating that all three MAPKs (extracellular signal-regulated kinase (ERK), p38, and c-Jun N-terminal kinase (JNK)) exhibit substrate specificity toward Cx43CT (34). In contrast, the phosphorylation status and kinase-mediated regulation of Cx47CT remain poorly understood.

The current study was designed to understand how RSA59 infection affects OLG cells and modulates oligodendrocytic Cx47-mediated GJ communication and LIF-LIFR/gp130 signaling, using a murine-β-coronavirus (RSA59)-induced in vivo model in combination with enriched primary OPC and OLG cultures. We further explored a potential link between these pathways by examining ERK, a downstream effector of LIF signaling, and its interaction with Cx47.

Here, we found that RSA59 targets oligodendroglial lineage cells including OPCs and mature OLGs, and elicits mature oligodendroglial apoptosis. RSA59 infection also reduced the expression of mature and myelinating OLG markers (CNPase and MBP) despite increased expression of the OPC marker (A2B5), suggesting impaired oligodendroglial maturation and myelination. Furthermore, RSA59 infection also alters key oligodendroglial survival and communication mechanisms mediated by Cx47 GJs and LIF-LIFR/gp130 signaling. Lastly, we found that ERK, a downstream effector of LIF signaling, phosphorylates the Cx47 C-terminal domain at the Serine-372 position, suggesting a potential regulatory mechanism for Cx47 function. Overall, RSA59-induced modulation of both Cx47-mediated oligodendroglial communication and the LIF-LIFR/gp130 signaling provides new mechanistic insights into chronic white matter pathology and identifies potential therapeutic targets for progressive demyelinating disorders, including MS. In addition, our findings also identify ERK-mediated phosphorylation of Cx47 as a potential regulatory mechanism and highlight it as a promising area for future investigation.

## Materials and Methods

### Isolation and Enrichment of Oligodendrocyte Precursor Cells (OPCs) from Neonatal Mouse Brain

Primary mixed glial cultures were prepared from postnatal day 0-1 mouse pups with minor modifications to previously described protocols (35). Following removal of the meninges, brain tissues were finely minced and incubated at 37°C for 30 min in a shaking water bath in Hanks’ Balanced Salt Solution (HBSS; GIBCO) containing 300 μg/ml DNase I and 10 mg/ml trypsin (Sigma). After enzymatic digestion, the dissociated cells were triturated in the presence of 0.25% fetal bovine serum (FBS), followed by washing and centrifugation at 300 × g for 10 min. The resulting cell pellet was resuspended in HBSS and passed through a 70 μm nylon mesh to remove debris. A second wash and centrifugation step (300 × g for 10 min) was then performed. Cells were subsequently diluted in initial plating medium consisting of Dulbecco’s essential medium supplemented with 1% penicillin-streptomycin, 1% non-essential amino acids, 0.2 mM L-glutamine, and 10% FBS, and maintained for 24 h at 37°C in a humidified CO₂ incubator. Non-adherent cells were removed after 24 h by washing with HBSS lacking Ca²⁺ and Mg²⁺, and the cultures were transitioned to serum-free growth medium composed of Neurobasal medium supplemented with B27, 10 ng/ml bFGF, 2 ng/ml PDGF, and 1 ng/ml NT-3. Cultures were maintained in this medium until confluence was reached (approximately 1 week). OPCs present within the mixed glial cultures, which primarily consisted of astrocytes and OPCs, were enriched using a wash-down method based on their differential adhesion properties. OPCs, which grow on top of the astrocytic layer and exhibit lower adherence compared to astrocytes, were selectively dislodged. Detached cells were collected and plated onto Poly-D-Lysine-coated coverslips and maintained in serum-free growth medium until approximately 80% confluence. The purity of the OPC cultures was assessed by double-label immunofluorescence using anti-A2B5 and anti-PDGFRα antibodies as OPC markers, along with nuclear counterstaining using 4′,6-diamidino-2-phenylindole (DAPI).

### Differentiation of Oligodendrocytes

Confluent OPC cultures were maintained in OLG differentiation medium consisting of DMEM: F12 (1:1) supplemented with 10% FBS, 10 μg/ml transferrin, 5 μg/ml insulin, and 30 nM sodium selenite for 7-10 days in accordance with previously established procedures. OPCs differentiated for 7 days (mature OLGs) were characterized using the mature oligodendrocyte-specific marker anti-CC1. In addition, differentiated OLG cultures were infected with RSA59 virus, followed by downstream experimental analyses.

### Infection of Primary OPCs and Differentiated OLGs with RSA59

RSA59, an isogenic recombinant strain derived from the parental neurotropic demyelinating mouse hepatitis virus strain MHV-A59, was used for all infection experiments as previously described (26, 36, 37). Primary OPCs were infected using infection medium consisting of Neurobasal medium supplemented with B27 and devoid of growth factors, containing RSA59 virus at a multiplicity of infection (MOI) of 1. Cells were incubated with the virus for 1.5 h at 37°C in a humidified CO₂ incubator. Following the 1.5 h incubation period, the infection medium was removed, and the infected cells were maintained in oligodendrocyte-specific growth medium composed of Neurobasal medium supplemented with B27 and the required growth factors (10 ng/ml bFGF, 2 ng/ml PDGF, and 1 ng/ml NT-3). Viral infectivity, assessed by viral EGFP expression, was monitored at 6, 12, 18, and 24 h post-infection (p.i.). The representative images were acquired using a 10× objective on a Nikon Eclipse Ts2-FL microscope (Tokyo, Japan) equipped with a Nikon DS-Fi3 camera (Tokyo, Japan) at respective time points. Similarly, mature oligodendrocyte cultures at day 7 post-plating (differentiation stage) were infected with RSA59 at an MOI of 1. At 24 h p.i., the culture medium was removed, and cells were fixed with 4% paraformaldehyde (PFA). Cells were subsequently processed for immunofluorescence staining using specific primary antibodies according to experimental requirements. Details of antibodies and dilutions are provided in Table 1.

**Table 1.** Primary antibodies used in this study.

| Sl. No. | Primary Antibody | Application | Dilution | Source |
| --- | --- | --- | --- | --- |
| 1 | Anti-Cx47 (polyclonal) | IF<br>WB | 1:500<br>1:1000 | Invitrogen, MA, USA |
| 2 | Mouse monoclonal anti-CC1 | IF | 1:400 | EMD Millipore, Co |
| 3 | Anti-PDGFRα | IF | 1:500 | Cell Signaling Technology (CST) |
| 4 | Anti-A2B5 | IF<br>WB | 1:5<br>1:10 | (Kind Gift from Judith B. Grinspan, Children's Hospital of Philadelphia, Philadelphia, PA) |
| 5 | Anti-Gal C | IF | 1:5 |  |
| 6 | Anti-PLP | IF | 1:10 |  |
| 7 | Anti-CNPase | WB | 1:10 |  |
| 8 | Anti-MBP | WB | 1:10 |  |
| 9 | Anti-pERK | WB | 1:1000 | D13.14.4E, 4370S, Lot no. 28 CST |
| 10 | Anti-ERK (Total) | WB | 1:1000 | 137F5, CST |
| 11 | Anti-β actin | WB | 1:10000 | Invitrogen, MA, USA |
| 12 | Anti-gp130 | WB | 1:1000 | Cell Signaling Technology (CST) |

### Immunofluorescence of Primary OPCs and Mature Oligodendrocytes

Immunofluorescence analysis was carried out as previously described, with minor modifications (27, 28). Primary OPCs and mature OLGs were seeded onto etched glass coverslips, followed by infection with RSA59 virus. At 24 h p.i., cells were fixed with 4% PFA for 10 min at room temperature. Cells were permeabilized using phosphate-buffered saline (PBS) containing 0.2% Triton X-100 and subsequently blocked in PBS supplemented with 1% bovine serum albumin (BSA). Incubation with primary antibodies diluted in blocking solution was performed overnight at 4°C. After washing, cells were incubated with the appropriate secondary antibodies diluted in blocking solution. Following this, cells were washed with PBS containing 0.2% Triton X-100 and mounted using mounting medium containing DAPI (VectaShield, Vector Laboratories). Slides were visualized and images were acquired using a Nikon Eclipse Ti2 inverted confocal microscope (Nikon Corp.) equipped with an AXR confocal scanning module and controlled by NIS-Elements ER software, using 60X or 100X oil immersion objectives with 1.5X zoom. Images used for representative purposes and quantification were captured at 100X and 60X magnification, respectively. Subsequent image processing and quantitative analyses were performed using ImageJ/Fiji (NIH, USA).

### Ethics approval

All animal experiments conducted in this study were approved by the Institutional Animal Ethics Committee (IAEC) of the Indian Institute of Science Education and Research (IISER) Kolkata and were carried out in strict accordance with the prescribed institutional guidelines. All experimental procedures were performed in accordance with the statutory rules and regulations laid down by the Committee for the Control and Supervision of Experiments on Animals (CCSEA), Government of India.

The protocols to obtain pups for primary neuroglial culture experiments were approved under IAEC Protocol numbers: **IISERK/IAEC/AP/2019/29.01** and **IISERK/IAEC/AP/2019/29.02.** Similarly, animal experiments involving adult mice were conducted under the validated approval of IAEC protocol numbers: **IISERK/IAEC/AP/2024/125** and **IISERK/IAEC/AP/2026/182**.

### Murine Coronavirus (RSA59) Infection of Mice

RSA59, an isogenic recombinant strain derived from the parental neurotropic demyelinating mouse hepatitis virus strain MHV-A59, was used for *in vivo* infection as previously described (38). Four-week-old male C57BL/6 mice were inoculated intracranially with 20,000 plaque-forming units (pfu) of RSA59, corresponding to the 50% LD₅₀ dose. Mock-infected control mice were inoculated intracranially with vehicle (PBS supplemented with 0.75% BSA) and were maintained in parallel. Following infection, mice were monitored daily for the development of clinical signs, including ruffled fur, reduced mobility, hind limb paralysis, weight loss, and mortality. Mice were euthanized at three defined disease stages: peak inflammation/acute phase (day 5 p.i.), acute-to-chronic transition phase (day 10 p.i.), and peak demyelination/chronic phase (day 30 p.i.). Brain tissues were subsequently harvested according to the specific experimental requirements. All animal experiments were conducted in strict accordance with institute ethical guidelines and applicable regulations.

### Tissue Processing, Immunofluorescence Staining on Mouse Brain Cryosections, and Microscopy

RSA59- and mock-infected mice at 5, 10, and 30 days p.i. were transcardially perfused with 1× PBS, followed by fixation with 4% paraformaldehyde (PFA) in 1× PBS. Brains were post-fixed in 4% PFA for 6 h and cryoprotected by sequential incubation in 10% sucrose in PBS for 4 h and 30% sucrose in PBS overnight at 4°C. Tissues were embedded in OCT compound and coronally sectioned at a thickness of 10 µm using a cryostat (Thermo Scientific). Sections were mounted onto charged glass slides. Immunostaining was performed as previously described (28). Cryosections were allowed to equilibrate to room temperature for 10 min and then treated with chilled ethanol for 10 min at −20°C. Sections were washed with 1× PBS and incubated in 1 M PBS-glycine for 1 h at room temperature, followed by treatment with 1 mg/ml NaBH₄ in PBS for 10 min to reduce autofluorescence. Permeabilization was carried out using 0.4% Triton X-100 for 20 min at room temperature. Blocking was performed with 1% BSA and 0.25% Triton X-100 in 1× PBS for 1 h at room temperature. Sections were incubated overnight at 4°C in a humidified chamber with primary antibodies diluted in antibody diluent containing 0.25% BSA and 0.1% Triton X-100 in 1× PBS. The following day, sections were washed and incubated for 2 h at room temperature with appropriate Alexa Fluor 488-, Alexa Fluor 568-, or Alexa Fluor 647-conjugated secondary antibodies (1:1000). All incubation steps were conducted in a humidified chamber. After washing, sections were mounted using mounting medium containing DAPI (VectaShield, Vector Laboratories). Confocal imaging was performed using a Leica SP8 confocal laser scanning microscope (Leica Microsystems) equipped with an HC PL APO ×63 oil immersion objective, or a Zeiss LSM710 confocal microscope. Excitation was achieved using 405-, 488-, 552-, and 647-nm laser lines. Image acquisition and processing were conducted using LAS X software (Leica Microsystems) or ZEN 3.7 software (Carl Zeiss AG), respectively. Quantitative image analysis was performed using ImageJ/FIJI software (NIH, USA).

### TUNEL Assay

Frozen brain sections (Coronal sections of 10-μm thickness) from mock- and RSA59-infected mice at day 5, day 10, and day 30 p.i. were used for combined tissue immunofluorescence (as described earlier in this manuscript) and subsequent TUNEL staining. Mature Oligodendrocytes were first stained with the specific marker anti-CC1 (dilution mentioned in Table 1), followed by incubation with the appropriate Secondary antibody. Thereafter, TUNEL staining was performed according to the manufacturer’s instructions (Roche Diagnostics, In Situ Cell Death Detection Kit, TMR red, Cat.no 12156792910) and validated by colocalization with DAPI-stained nuclei. Images were captured using a Leica confocal microscope. Further, the acquired images were processed and quantified using ImageJ/Fiji software.

### Protein Isolation and Immunoblot Analysis

Protein isolation was performed according to a previously described protocol with minor modifications (39). Brain tissue was collected from euthanized mice following transcardial perfusion with 20 mL PBS and flash-frozen in liquid nitrogen. Tissues were lysed in RIPA buffer containing 0.1% SDS and 0.1% Triton X-100, supplemented with 1× Complete Mini protease inhibitor cocktail tablets (#11836153001; Roche) and phosphatase inhibitors (10 mM sodium orthovanadate, 10 mM sodium fluoride, and 10 mM sodium pyrophosphate). Brain tissues were homogenized by trituration followed by sonication. Lysates were centrifuged at 13,500 rpm for 30 min at 4°C, and the supernatant was collected as the total protein extract. Protein concentration was determined using the Pierce BCA protein assay kit (Thermo Scientific, Rockford, IL, USA). Equal amounts of protein were resolved on 10% SDS-PAGE gels and transferred onto PVDF membranes. Membranes were blocked with 5% non-fat skim milk prepared in TBST (Tris-buffered saline containing 0.1% Tween-20) for 1 h at room temperature, followed by overnight incubation at 4°C with primary antibodies including anti-Cx47, anti-pERK, anti-A2B5, anti-CNPase, or anti-MBP at the dilutions specified in Table 1. Membranes were subsequently washed with TBST and incubated with appropriate HRP-conjugated secondary antibodies (1:10,000). Immunoreactive bands were detected using the SuperSignal™ West Pico PLUS Chemiluminescent Substrate (Thermo Fisher Scientific) and visualized using the Syngene G:BOX ChemiDoc system with GENESys software (Genesys, Cambridge, UK). To verify equal protein loading, all membranes were reprobed with anti-β-actin antibody. Membranes probed for pERK were additionally reprobed with anti-total ERK antibody. Densitometric analysis of immunoblots were performed using ImageJ software (NIH, USA).

### Quantification of Mouse LIF (mLIF) in primary neuroglial cell culture supernatant by ELISA

The murine primary astrocytes, oligodendrocytes, and neurons were cultured separately in a 60mm dishes following a previously established procedure (35). Once the cells were 70-80% confluent, infected with the RSA59 virus at MOI 1 for 24 hr in low serum-containing medium. After 24 hours, the culture supernatant was collected from the RSA59-infected and mock-infected cultures. Secretory mouse LIF (mLIF) protein levels in the culture media were assessed using the Mouse LIF DuoSet ELISA kit (Cat No: DY449; R&D Systems, Minneapolis, MN, USA) following the manufacturer’s instructions.

### Gene Expression Analysis: RNA Isolation, Reverse Transcription, and Quantitative Polymerase Chain Reaction

Total RNA was isolated from flash-frozen whole brain tissues of RSA59- and mock-infected mice following transcardial perfusion with 1X PBS, using the TRIzol extraction method. RNA concentration and purity were determined using a NanoDrop ND-2000 spectrophotometer. For complementary DNA (cDNA) synthesis, 1 µg of total RNA was reverse-transcribed using the High-Capacity cDNA Reverse Transcription Kit (Applied Biosystems) according to the manufacturer’s instructions. Quantitative real-time PCR (qRT-PCR) was performed using the iTaq Universal SYBR Green qPCR kit (Bio-Rad) on a Bio-Rad CFX Real-Time PCR system. Amplification was carried out under the following cycling conditions: initial denaturation at 95°C for 7 min, followed by 40 cycles of denaturation at 95°C for 10 s and annealing/extension at 60°C for 30 s. A melting curve analysis was performed at 60°C for 30 s to confirm amplification specificity. All reactions were conducted in quadruplicate. Relative gene expression was calculated using the comparative threshold (ΔΔCt) method. Target gene expression levels in RSA59- and mock-infected mice were normalized to GAPDH and expressed as fold change relative to the corresponding mock-infected controls. Primer sequences are provided in Table 2.

**Table 2.**
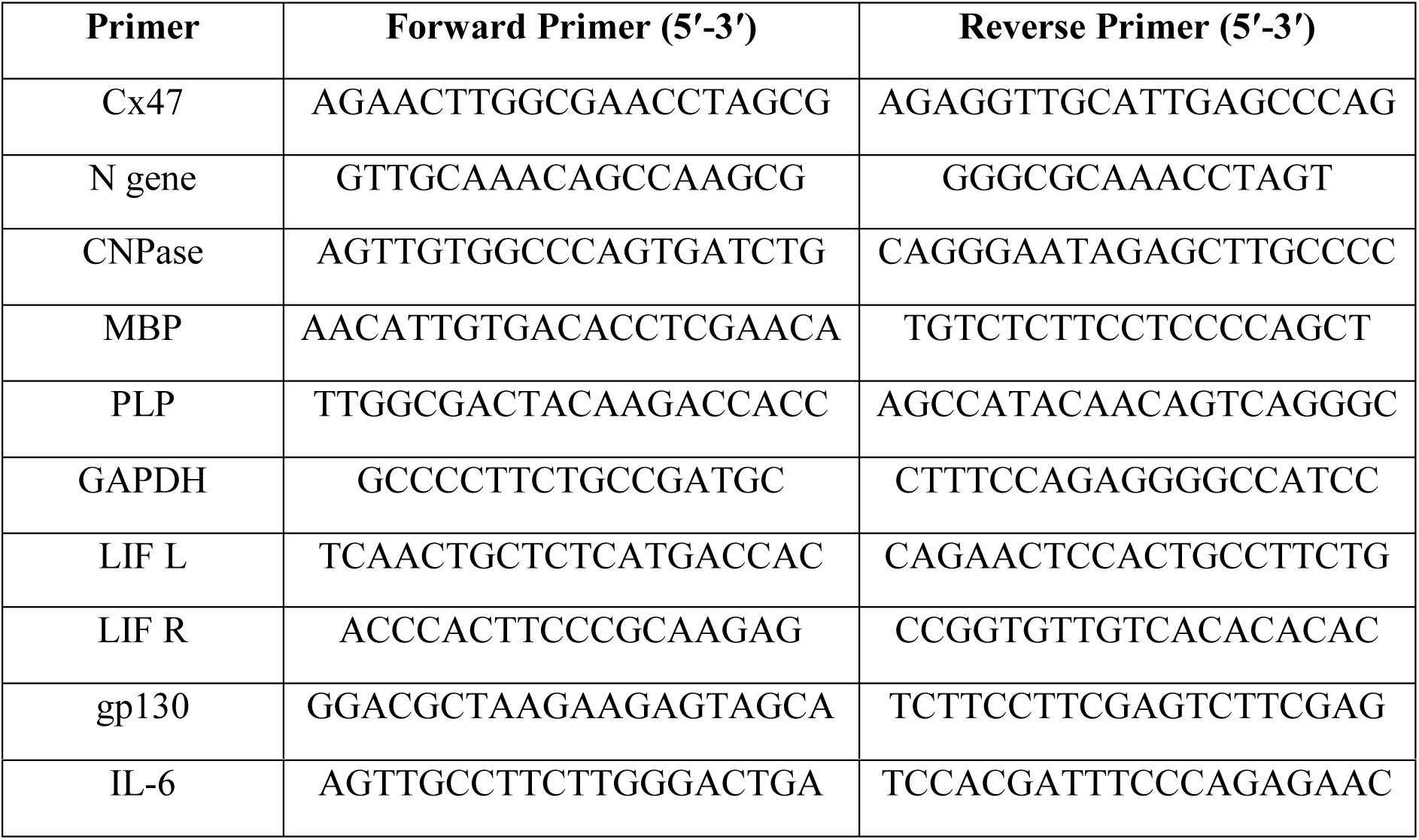
Sequences of primers used in the study.

### Quantification of Fluorescence Images

Coronal brain Cryosections from RSA59- and mock-infected mice were collected at 5, 10, and 30 days p.i. (n=4-6 animals per group). Sections were triple-immunolabeled with anti-PLP, anti-Cx47, and anti-CC1 antibodies at the dilutions listed in Table 1 to identify mature/myelinating oligodendrocytes. Double immunolabeling with anti-A2B5 and anti-Cx47 antibodies was performed to identify OPCs, as described above. Mean fluorescence intensity (MFI) of Cx47 puncta in PLP⁺/CC1⁺ and A2B5⁺ cells was determined from confocal image stacks by merging all optical sections using ImageJ software. For each PLP⁺/CC1⁺ cell, a region of interest (ROI) was manually drawn using the freehand selection tool. Similarly, ROIs were defined for individual A2B5⁺ cells. The MFI of Cx47 within the selected ROIs (OPCs and OLGs) was then calculated using ImageJ/Fiji software. Cx47 MFI values were normalized to the corresponding mock-infected controls and expressed as relative changes in MFI in PLP⁺/CC1⁺ and A2B5⁺ cells (Figure 4-6). *In vitro* Cx47 MFI was quantified in a similar manner by specifically marking CC1⁺ mature OLGs and A2B5⁺ OPCs (Figure 7). *In vivo* RSA59 EGFP expression and *in vitro* RSA59 viral kinetics were quantified by measuring EGFP fluorescence intensity using ImageJ/Fiji software. Absolute values were plotted as the percentage area of viral EGFP expression. All quantifications were performed in a blinded and randomized manner.

### Coupled Kinase Assay

Activity of ERK2 kinase was measured by monitoring the rate of phosphorylation of 74µg MBP bovine protein (Thermo, #cat number: 13228010) and 131µg CX47 Peptide (DRDSPPCAGLNAT). Phosphorylation was monitored by measuring the oxidation of NADH to NAD+. The reaction buffer consists of 25mM HEPES (pH 7.5), 100mM NaCl, 20mM Mgcl2, 1.4mM phosphoenolpyruvate (PEP), 1mg/ml NADH, 75 U/ml pyruvate kinase, 105 U/ml lactate dehydrogenase and 5mM ATP. The reaction was initiated by the addition of 6µg of activated ERK2 and a decrease in absorbance at 340nm was monitored for 120 minutes at 25 °C in SpectraMax M2e plate reader. The specific activity of ERK2 toward MBP and the CX47 peptide was determined by calculating the ratio of the reaction velocity (µmol/min), obtained from the slope of the decrease in absorbance to the total amount of ERK2 used in the reaction.

### ERK expression and Purification

A co-expressing MEK1-ERK2 plasmid was obtained from Addgene (#39212) and expressed in BL21 (DE3). The bacterial cells were grown in 37 °C to an O.D600 of 1.4 before inducing with 0.25mM IPTG for 14 hour at 30 °C. Before purification, the bacterial cells were harvested in Buffer A (25mM HEPES (pH 7.5), 300mM NaCl, 10mM imidazole, 5% glycerol). Cells were treated with 1mg/ml Lysozyme for 1 hour and then, sonicated. After centrifugation in 14000 RPM, the soup was loaded in Nickel column (Cytiva) and eluted with Buffer B (25mM HEPES (pH 7.5), 300mM NaCl, 750mM imidazole, 5% glycerol). The eluted protein was diluted in Buffer QA (25mM HEPES (pH 7.5), 1mM β-mercaptoethanol, 5% glycerol) and loaded into anion exchange column (Cytiva-Q). Then, the protein was eluted by gradient of Buffer QB (25mM HEPES (pH 7.5), 1M NaCl, 1mM β-mercaptoethanol, 5% glycerol). Selected pure fractions were concentrated and loaded into Sepharose gel filtration column and finally stored in storage buffer (25mM HEPES (pH 7.5), 150mM NaCl, 1mM β-mercaptoethanol, 10% glycerol). Thereafter, purified ERK2 was concentrated, aliquoted, flash freezed in liquid nitrogen and stored at −80 °C.

### Statistical Analysis

All experimental data were expressed as mean ± standard error of the mean (SEM). Statistical comparisons between groups were performed using a two-tailed Student’s *t*-test with Welch’s correction. A *P* value of < 0.05 was considered statistically significant (ns - not significant, *P < 0.05, **P < 0.01, ***P < 0.001, ****P < 0.0001). All statistical analyses were conducted using GraphPad Prism 8 software (GraphPad Software, Inc., La Jolla, CA, USA).

## Results

### 1. Murine Coronavirus (MCoV) RSA59 exhibits Oligodendroglial Tropism both *in vitro*and *in vivo*

Previous studies have demonstrated that Mouse Hepatitis Virus-A59 (MHV-A59) infects CNS glial cells, including astrocytes and oligodendrocytes, impairing their gap junction intercellular communication and resulting in virus-induced chronic progressive demyelination (27, 28, 37). Therefore, before proceeding with further detailed experiments, we aimed to evaluate the oligodendroglial tropism of RSA59, an isogenic recombinant strain derived from the dual hepato-neurotropic strain MHV-A59. We employed complementary in vitro and in vivo approaches. *In vitro* studies were performed using primary OPCs isolated from neonatal (P0) C57BL/6 mouse pups, following a previously published protocol (35). After isolation, OPCs were infected with RSA59 at a multiplicity of infection (MOI) of 1. Viral infectivity was monitored by EGFP expression at 6 h, 12 h, and 18 h p.i. (Fig. S1) and at 24 h p.i. (Fig. 1A, B). Immunofluorescence studies demonstrated colocalization of RSA59-EGFP with the OPC marker A2B5, suggesting specific tropism toward oligodendrocyte lineage cells (Fig. 1C). Viral infectivity kinetics were further assessed by quantifying EGFP expression, which showed a progressive and significant increase from 6 h to 24 h p.i. *in vitro* (Fig. 1D). Similarly, the absolute number of RSA59-EGFP-positive OPCs also significantly increased from 6 h to 24 h p.i. (Fig. 1E).

**Figure 1.**
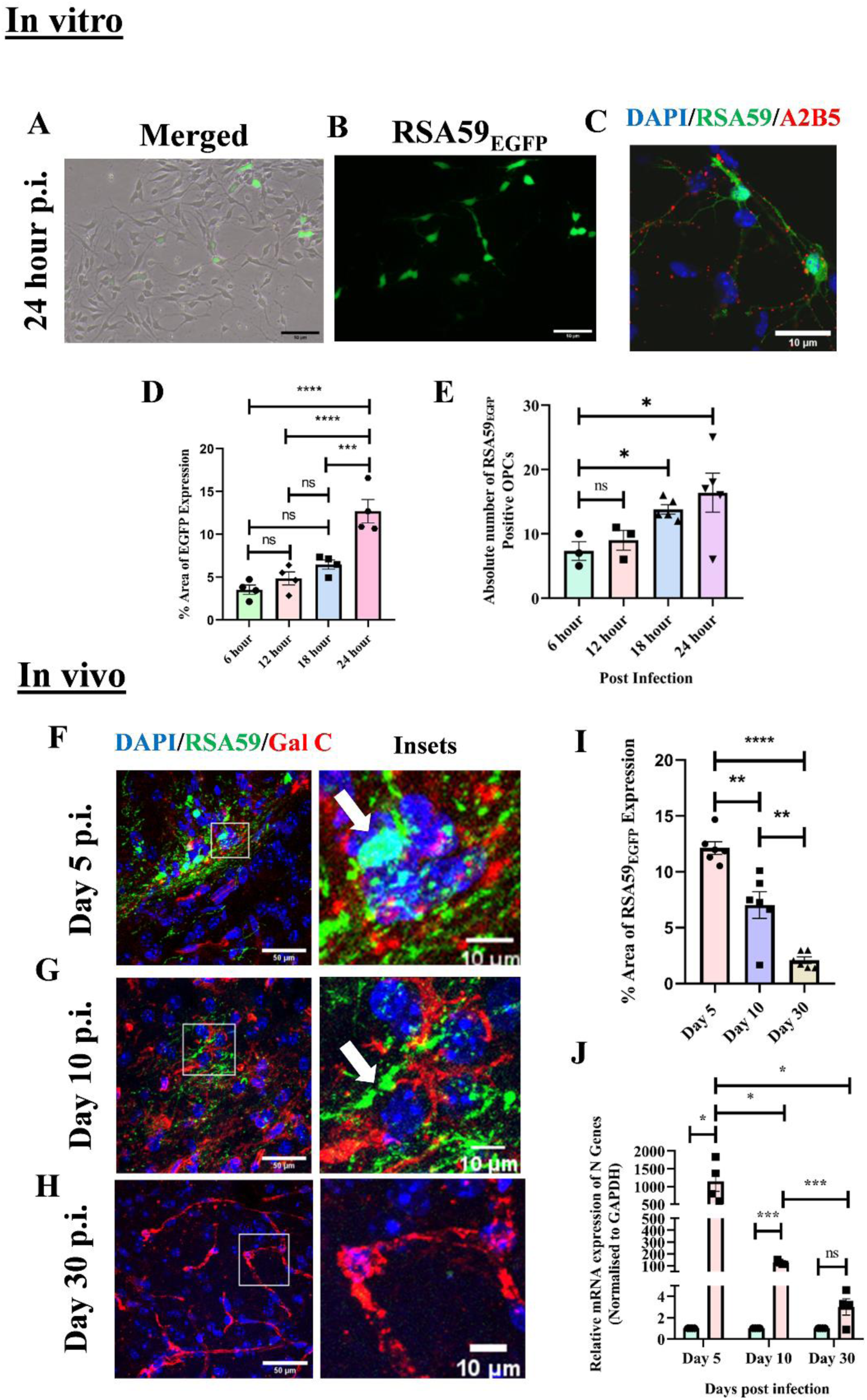
RSA59 infects oligodendroglial lineage cells both *in vitro* and *in vivo*. (A, B) Representative bright-field and EGFP staining photomicrographs of murine primary OPCs infected with RSA59 at MOI 1 for 24 h p.i. (C) Representative confocal images of OPCs marked by A2B5 (red) and infected with RSA59 EGFP (green). (D) Quantification of virus (RSA59) EGFP spread (% of area) and (E) absolute number of RSA59-infected (RSA59+) OPCs from 6 h p.i. to 24 h p.i. (F-H) Representative confocal images of coronal brain sections stained with mature oligodendrocyte marker (Gal C) and RSA59 EGFP (green) at day 5 p.i., day 10 p.i., and day 30 p.i., respectively. The boxed areas are shown as manually zoomed insets just adjacent to the main image. The white arrow indicates infected oligodendrocytes. Nuclei were counterstained with DAPI. (I) Quantification of EGFP expression in the brain tissues at day 5 p.i., day 10 p.i., and day 30 p.i. (J) Viral N gene mRNA expression quantification in d5, d10, and d30 post RSA59-infected brain tissues. The results are expressed as mean ± SEM (*in vivo*: n = 4-6 animals/per group, *in vitro*: n=3 per group). Asterisks represent statistical significance calculated using unpaired Student’s t-test and Welch correction. P < 0.05 was considered significant. ns - not significant, *P < 0.05, **P < 0.01, ***P < 0.001, ****P < 0.0001.

In vivo, intracranial RSA59 infection of C57BL/6 male mice followed three characteristic disease stages: an acute neuroinflammatory stage (day 5 p.i.), an acute-chronic transitional phase (day 10 p.i.), and finally a chronic demyelinating phase (day 30 p.i.), as described previously (38). Accordingly, these three time points were evaluated for viral EGFP expression. Immunofluorescence on brain sections at these time points revealed colocalization of the mature OLG marker Galactocerebroside C (Gal C) with RSA59-EGFP, confirming that RSA59 targets oligodendrocytes *in vivo* (Fig. 1F & G; Insets). A gradual clearance of viral EGFP expression was observed, with no detectable expression at day 30 p.i. (Fig. 1H; Insets). Quantification showed a significant decrease in the expression of RSA59-EGFP from day 5 p.i. to day 30 p.i. in vivo (Fig. 1I). However, viral N gene mRNA transcripts persisted during all the stages of RSA59 infection and were significantly upregulated at day 5 and day 10 p.i., with an increasing trend still detectable at day 30 p.i. (Fig. 1F). Collectively, these findings demonstrate that the neurotropic RSA59 virus directly targets oligodendrocyte lineage cells including OPCs and mature OLGs, both *in vitro* and *in vivo*. These observations prompted further investigation into the impact of RSA59 infection on oligodendrocytic gap junction (GJ) protein Cx47 and the LIF-LIFR/gp130 signaling axis.

### 2. RSA59 infection triggers mature oligodendroglial apoptosis in the brain

Following the observation that RSA59 infects oligodendroglial lineage cells (OPCs and OLGs), we next sought to decipher its consequences, as viral infections have previously been shown to trigger direct cytopathic damage as well as apoptotic death. Therefore, we further examined whether Murine Coronavirus (RSA59) infection also induces apoptotic cell death. We performed the colocalization of mature oligodendrocyte marker CC1 with TUNEL (a dye that specifically binds to fragmented DNA of apoptotic cells) in the brain tissue during acute (d5), acute-chronic (d10), and chronic (d30) stages of RSA59 infection.

The result showed the presence of apoptotic mature OLGs during all the stages post RSA59 infection (Fig. 2A, B & C, and Insets). However, mature OLGs apoptosis is significantly elevated during the acute neuroinflammatory (d5 p.i.) stage and gradually decreases towards d10 and d30 p.i. (Fig. 2D). In contrast, no colocalization of CC1 and TUNEL was observed in mock-infected brain tissues across all the examined stages of RSA59 infection (data not shown). Collectively, these results indicate that RSA59-induced oligodendroglial apoptosis during the early phase (d5 and d10) of infection might be one of the contributory factors leading to subsequent chronic, progressive demyelination that peaks at day 30 p.i.

**Figure 2.**
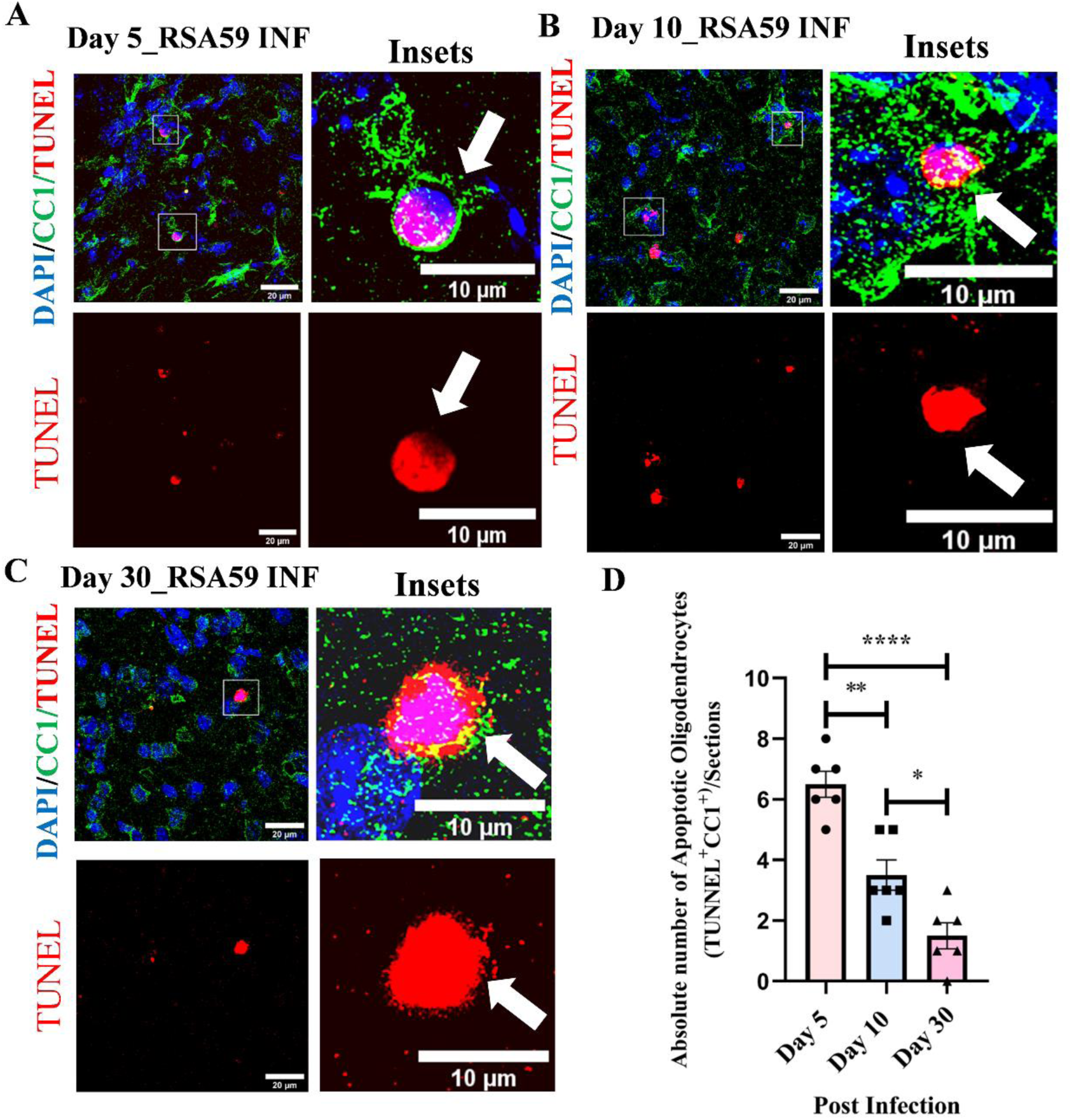
RSA59 infection induces apoptosis of mature oligodendrocytes in the brain. (A-C) Representative confocal images of brain sections showing mature oligodendrocytes marked with CC1 (green) and apoptotic cells detected by TUNEL (red), with merged images showing CC^+^/TUNEL^+^ cells (A) during the acute stage (d5 p.i), (B) acute-chronic transition stage (d10 p.i.), and (C) the chronic stage (d 30 p.i) of RSA59 infection. The boxed areas are shown as mannualy zoomed insets just adjacent to the main image. White arrows show apoptotic mature OLGs marked with both CC1 and TUNEL. Nuclei are counterstained with DAPI. (D) Scatter plot showing Quantification of apoptotic mature OLGs (TUNEL+ and CC1+) per field during different stages post RSA59 infection. The results are expressed as mean ± SEM (n = 4-6 animals/per group). Asterisks represent statistical significance calculated using unpaired Student’s t-test and Welch correction. P < 0.05 was considered significant. ns - not significant, *P < 0.05, **P < 0.01, ***P < 0.001, ****P < 0.0001

### 3. RSA59 infection differentially regulates the expression levels of OPCs, mature and myelinating oligodendrocyte cell-specific markers in the brain

In addition to inducing apoptosis, virus infection may also modulate the expression of various oligodendroglial cell-specific proteins essential for their structural integrity and function. To determine whether RSA59 infection in the brain alters the steady-state expression of oligodendroglial lineage markers during three distinct stages of RSA59 infection. We evaluated the protein levels of oligodendroglial lineage markers, including A2B5 (for OPCs), CNPase (for mature oligodendrocytes), and MBP (for myelinating oligodendrocytes), by immunoblotting across all stages of infection.

RSA59 infection persistently downregulated CNPase and MBP protein expression in the brain during the acute stage (day 5 p.i.; Fig. 3A), the acute-chronic transition stage (day 10 p.i; Fig. 3B), and the chronic stage (day 30 p.i.; Fig. 3C) compared with respective mock-infected controls. This sustained reduction in myelin-associated proteins is consistent with the progressive demyelination that characterizes chronic RSA59 infection as reported previously. In contrast, a significant increase in the OPC marker A2B5 expression was observed at all the stages (day 5, day 10, and day 30) following RSA59 infection, compared with mock-infected controls (Fig. 3 A-C). To further examine transcriptional regulation, mRNA transcript levels of mature and myelinating OLG markers (CNPase, MBP, and PLP) were analyzed. Differential expression was observed across distinct stages of RSA59 infection. MBP mRNA expression was significantly downregulated during the acute-chronic transition phase (day 10 p.i.; Fig. 3D). In contrast, PLP and CNPase mRNA levels were significantly reduced during the acute (day 5 p.i.) and chronic (day 30 p.i.) stages (Fig. 3E and 3F), while no significant differences were detected during the other stages for these specific markers. Collectively, the results demonstrate that RSA59 infection differentially regulates both mRNA and protein expression levels of mature and myelinating OLG markers across the distinct stages of RSA59 infection.

**Figure 3.**
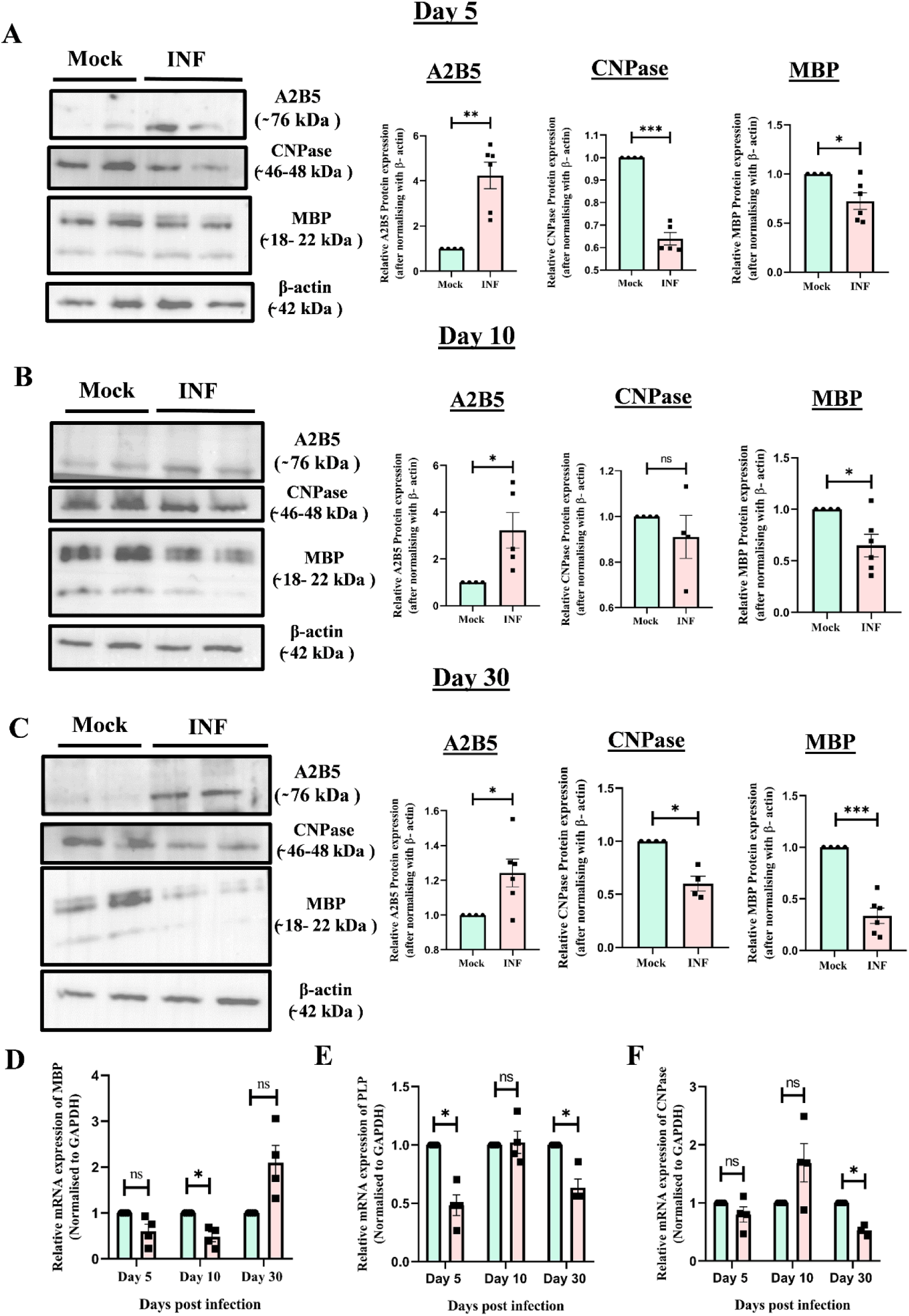
Differential expression of oligodendroglial lineage marker levels post RSA59 infection in the brain. (A-C) Representative immunoblots and corresponding scatter plots are depicting protein expression of oligodendroglial lineage cell (OPCs and OLGs) specific markers A2B5 (̴ 76 KDa), CNPase (2’3’-cyclic nucleotide 3’-phosphodiesterase, (46-48KDa) and MBP (myelin basic protein 18.5-21.5 KDa) in the brain of mock and RSA59 infected mice (A) at day 5 p.i, (B) day 10 p.i. and (C) day 30 p.i. Note: OPCs marker (A2B5) protein expression levels significantly increase in infected groups compared to the representative control groups. However, a persistent downregulation of mature and myelinating oligodendrocytes marker (CNPase & MBP) protein levels in infected samples compared to control during acute, acute-chronic and chronic phase of RSA59 infection. (D-F) Scatterplot of differential {(D) MBP, (E) PLP and (F) CNPase} mRNA expression in mock and RSA59-infected mouse brain post d5, d10 and d30 p.i. as measured by qRT-PCR. The results were expressed as mean ± SEM (n = 4-6 animals/per group). Asterisks represent statistical significance calculated using unpaired Student’s t-test and Welch correction. P < 0.05 was considered significant. ns - snot significant, *P < 0.05, **P < 0.01, ***P < 0.001, ****P < 0.0001.

Overall, the findings indicate that the persistent downregulation of mature and myelinating OLGs may be attributed to apoptosis of mature OLGs or vice versa. Whereas the sustained elevation of the OPC marker A2B5 at day 5, day 10 and day 30 p.i. implies an endogenous reparative response to restore oligodendroglial homeostasis following RSA59-induced damage.

### 4. Differential Modulation of Oligodendroglial GJ Protein Cx47 Expression in the Brain following RSA59 Infection

To evaluate the impact of RSA59 infection on oligodendrocytic GJ protein Cx47 expression, we assessed both total protein expression by immunoblotting and cell-type-specific Cx47 mean fluorescence intensity (MFI) in brain tissues at day 5 p.i (acute phase), day 10 p.i. (acute-chronic transition phase) and day 30 p.i. (chronic phase). Cx47 mean fluorescence intensity (MFI) was evaluated specifically in PLP^+^/CC1^+^ mature/myelinating oligodendrocytes and A2B5+ oligodendrocyte precursor cells (OPCs). Immunolabeling studies showed the presence of Cx47 puncta, representing oligodendrocytic GJs, on the soma and processes of PLP+/CC1+ mature/myelinating oligodendrocytes. Similarly, Cx47 puncta were observed in A2B5+ OPCs (Fig. 4-6).

**Figure 4.**
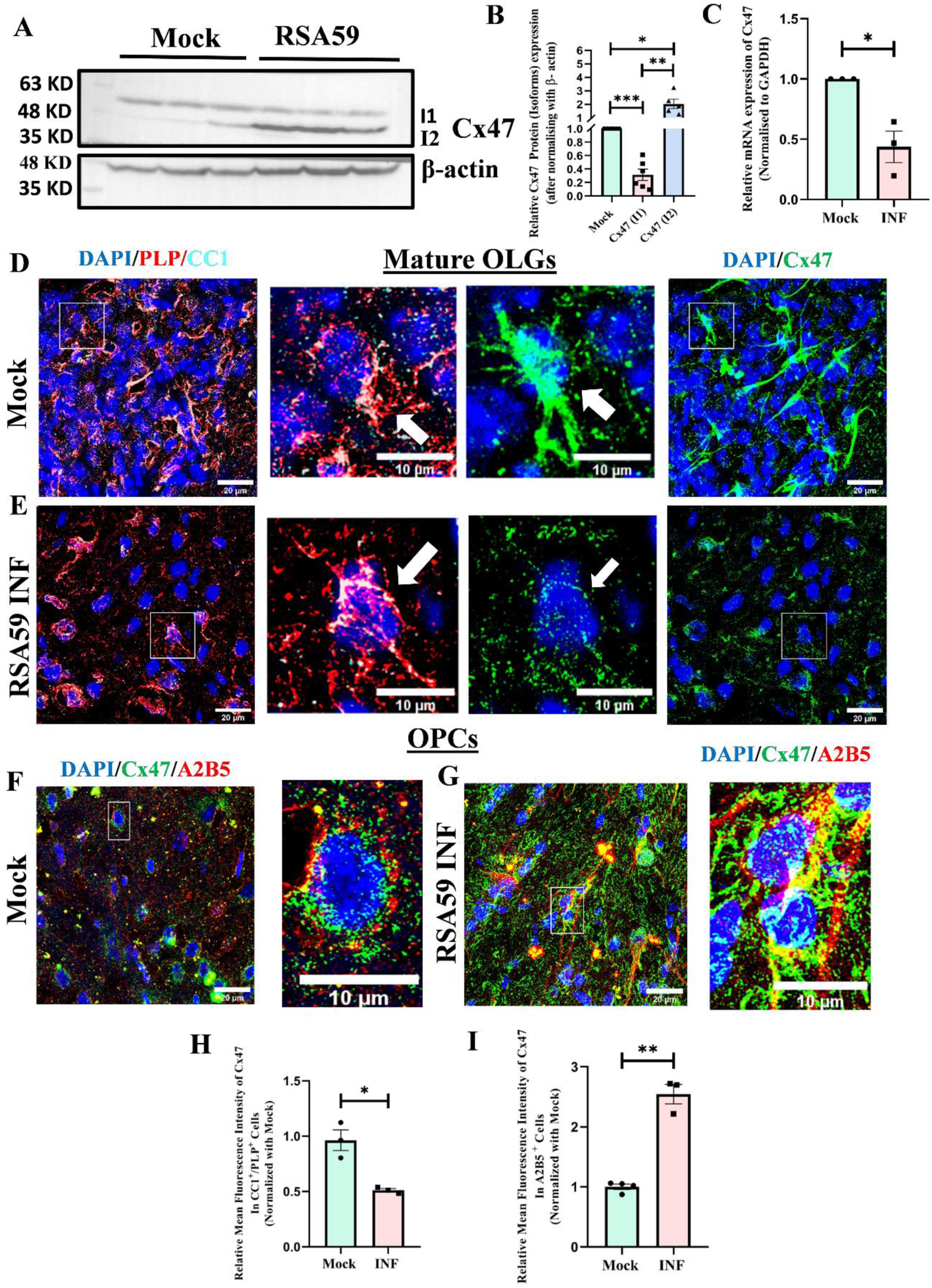
RSA59 differentially modulates Cx47 expression in the brain during the acute stage (d5 p.i.) of infection. (A) Representative Immunoblots and (B) corresponding scatterplot of two isoforms (I1=upper band, phosphorylated isoform; lower band: I2=non-phosphorylated isoform) of Cx47 protein isolated at d5 p.i. from RSA59-infected and mock-infected mouse brain tissues. (C) Scatter plot of Cx47 mRNA expression during the acute phase (d5 p.i.) in mock- and RSA59-infected mouse brain measured by quantitative PCR. (D) Mean fluorescence intensity (MFI) expression of Cx47 in the brain of mock-infected and (E) RSA59-infected (d5) mice was analysed by triple immunolabeling with anti-PLP (red), anti-Cx47 (green) and anti-CC1 (cyan) followed by confocal microscopy. Boxed regions are represented as insets, just adjacent to the main images. White arrows indicate specific Cx47 expression in both PLP+ and CC1+mature OLGs. (F) Expression of OPC-specific Cx47 in the brain of mock-infected and (G) RSA59-infected mice was analysed by immunolabeling with A2B5 (red) and anti-Cx47 (green), followed by confocal microscopy. Boxed regions are represented as insets, just adjacent to the main images (H & I). Scatterplots represent the quantification of mean fluorescence intensity (MFI) of Cx47 in PLP^+^ CC1^+^ mature/myelinating oligodendrocytes and A2B5^+^ OPC. Results were expressed as mean ± SEM (n=4-6 animals/per group). Asterisks represent statistical significance calculated using unpaired Student’s t-test with Welch correction. P < 0.05 was considered significant. ns - not significant, *P < 0.05, **P < 0.01, ***P < 0.001, ****P < 0.0001

**Figure 5.**
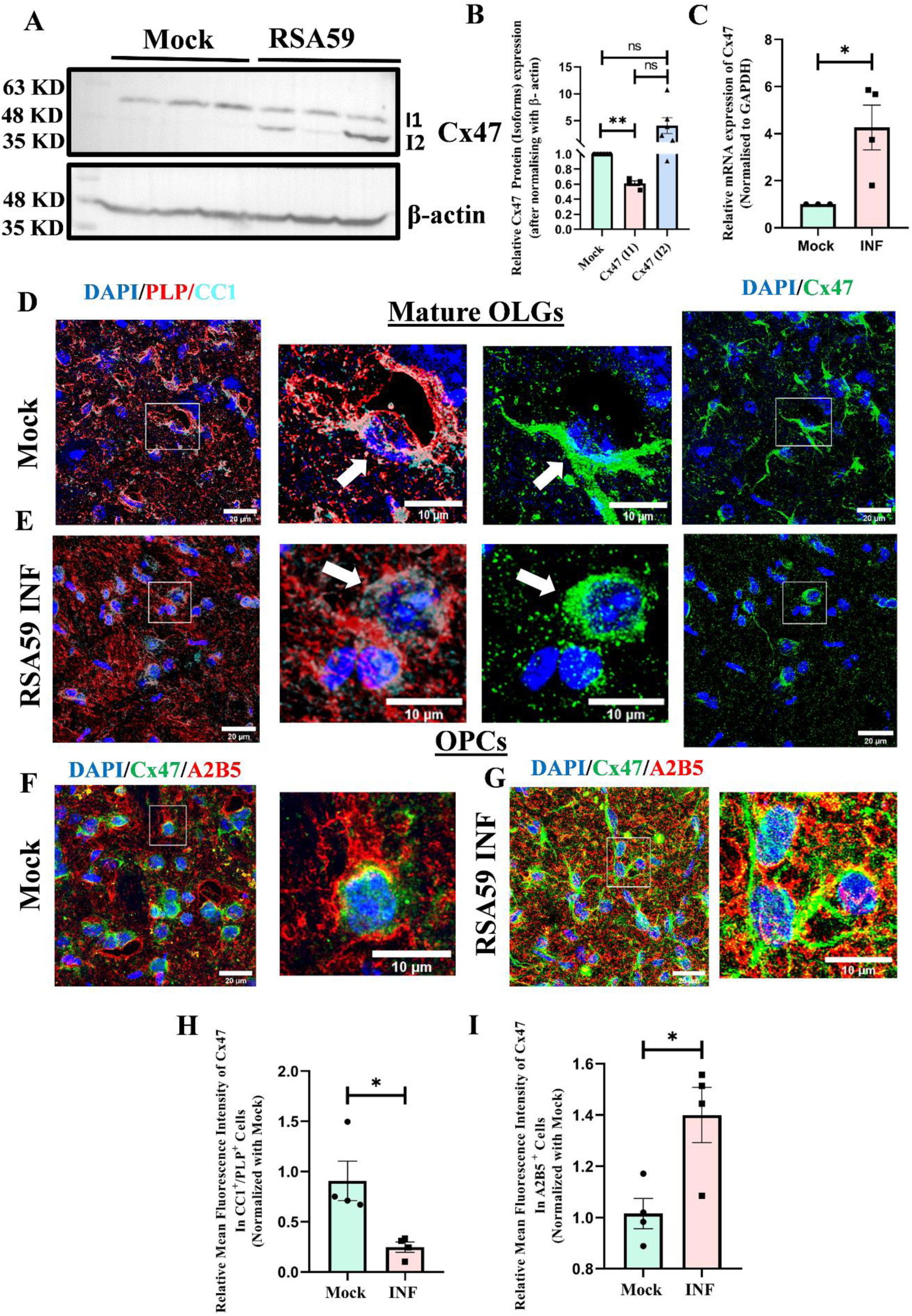
RSA59 differentially modulates Cx47 expression in the brain during acute-chronic transition stage (d10 p.i.) of RSA59 infection. (A) Representative Immunoblots and (B) corresponding scatterplot of two isoforms (I1=upper band, phosphorylated isoform; lower band: I2=non-phosphorylated isoform) of Cx47 protein isolated at d5 p.i. from RSA59-infected and mock-infected mouse brain tissues. (C) Scatter plot of Cx47 mRNA expression during the acute phase (d5 p.i.) in mock- and RSA59-infected mouse brain measured by quantitative PCR. (D) Mean fluorescence intensity (MFI) expression of Cx47 in the brain of mock-infected and (E) RSA59-infected (d5) mice was analysed by triple immunolabeling with anti-PLP (red), anti-Cx47 (green) and anti-CC1 (cyan) followed by confocal microscopy. Boxed regions are represented as insets, just adjacent to the main images. White arrows indicate specific Cx47 expression in both PLP+ and CC1+mature OLGs. (F) Expression of OPC-specific Cx47 in the brain of mock-infected and (G) RSA59-infected mice was analysed by immunolabeling with A2B5 (red) and anti-Cx47 (green), followed by confocal microscopy. Boxed regions are represented as insets, just adjacent to the main images (H & I). Scatterplots represent the quantification of mean fluorescence intensity (MFI) of Cx47 in PLP^+^ CC1^+^ mature/myelinating oligodendrocytes and A2B5^+^ OPC. Results were expressed as mean ± SEM (n=4-6 animals/per group). Asterisks represent statistical significance calculated using unpaired Student’s t-test with Welch correction. P < 0.05 was considered significant. ns - not significant, *P < 0.05, **P < 0.01, ***P < 0.001, ****P < 0.0001.

**Figure 6.**
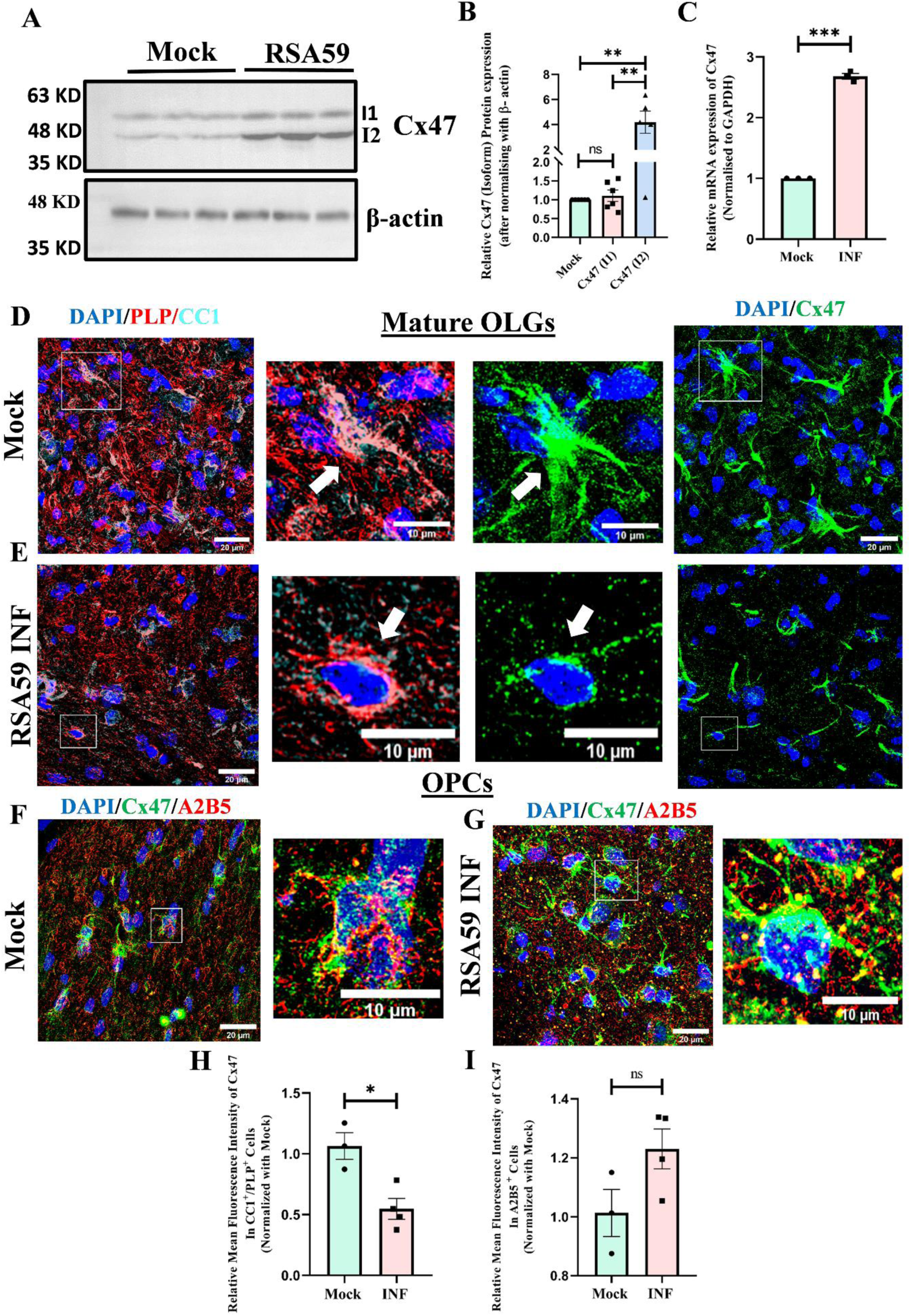
RSA59 differentially modulates Cx47 expression in the brain during chronic stage (d30 p.i.) of RSA59 infection. (A) Representative Immunoblots and (B) corresponding scatterplot of two isoforms (I1=upper band, phosphorylated isoform; lower band: I2=non-phosphorylated isoform) of Cx47 protein isolated at d5 p.i. from RSA59-infected and mock-infected mouse brain tissues. (C) Scatter plot of Cx47 mRNA expression during the acute phase (d5 p.i.) in mock- and RSA59-infected mouse brain measured by quantitative PCR. (D) Mean fluorescence intensity (MFI) expression of Cx47 in the brain of mock-infected and (E) RSA59-infected (d5) mice was analysed by triple immunolabeling with anti-PLP (red), anti-Cx47 (green) and anti-CC1 (cyan) followed by confocal microscopy. Boxed regions are represented as insets, just adjacent to the main images. White arrows indicate specific Cx47 expression in both PLP+ and CC1+mature OLGs. (F) Expression of OPC-specific Cx47 in the brain of mock-infected and (G) RSA59-infected mice was analysed by immunolabeling with A2B5 (red) and anti-Cx47 (green), followed by confocal microscopy. Boxed regions are represented as insets, just adjacent to the main images (H & I). Scatterplots represent the quantification of mean fluorescence intensity (MFI) of Cx47 in PLP^+^ CC1^+^ mature/myelinating oligodendrocytes and A2B5^+^ OPC. Results were expressed as mean ± SEM (n=4-6 animals/per group). Asterisks represent statistical significance calculated using unpaired Student’s t-test with Welch correction. P < 0.05 was considered significant. ns - not significant, *P < 0.05, **P < 0.01, ***P < 0.001, ****P < 0.0001.

Immunoblot analysis revealed that RSA59 infection dynamically regulated two distinct isoforms of Cx47 {Isoform 1 (I1) and Isoform 2 (I2)}. Cx47 resolved as a closely migrating doublet of bands, which were quantified separately at day 5 (Fig 4A), day 10 (Fig 5A), and day 30 p.i. (Fig. 6A). These closely migrating bands likely represent differential post-translational modifications of Cx47, presumably Cx47 phosphorylation. The upper band representing I1 (putative phosphorylated isoform) of Cx47 was reduced following RSA59 infection, whereas the lower band depicting I2 (putative non-phosphorylated) was significantly upregulated compared with mock-infected controls, indicating differential modulation of Cx47 isoforms by RSA59 infection. This differential isoform expression pattern of Cx47 was consistently observed during the acute phase (day 5 p.i.; Fig. 4B) and acute-chronic transition phase (day 10 p.i.; Fig. 5B). During the chronic phase (day 30 p.i.), I1 levels in RSA59-infected mice were comparable to mock-infected controls, whereas I2 remained significantly upregulated (Fig. 6B). Similar double-banding patterns and differential Cx47 regulation have previously been reported following infection with MHV-A59, the parental strain of RSA59 (30, 32, 40). Differential Cx47 mRNA expression was also observed across the three stages of RSA59 infection. During the acute stage (day 5 p.i.), mRNA levels were significantly downregulated (Fig. 4C). In contrast, significant upregulation was detected at day 10 p.i. and day 30 p.i. compared with mock-infected mice (Fig. 5C and 6C). The increased mRNA expression at later stages, together with the persistent downregulation of phosphorylated isoforms (I1) and upregulation of the non-phosphorylated isoforms (I2), suggests post-translational (likely phosphorylation-dependent) regulation of Cx47 following RSA59 infection.

Consistent with immunoblot findings, immunofluorescence analyses using anti-Cx47 in combination with anti-PLP and anti-CC1 antibodies showed a significant and persistent loss of Cx47 MFI (MFI/cell/field) in double marked PLP^+^/CC1^+^ mature/myelinating oligodendrocytes across all stages of RSA59 infection (day 5: Fig. 4, D & E; day 10: Fig. 5, D & E; day 30: 6 D & E; quantifications shown in Fig. 4H, 5H, and 6H). In contrast, a significant increase in Cx47 MFI (MFI/cell/field) was observed in A2B5^+^ OPCs at day 5 and day 10 p.i., with a non-significant increasing trend noted at day 30 p.i. (day 5: Fig. 4 F & G; day 10: Fig. 5 F & G; day 30: 6 F & G; quantifications shown in Fig. 4I, 5I, and 6I).

Thus, the differential expression of Cx47 isoforms in the brain upon RSA59 infection may be attributed to differential Cx47 expression in two different subpopulations of oligodendrocyte lineage cells (OPCs and OLGs) in the brain. PLP+/CC1+cells in mock-infected brain sections (Insets, White arrow) displayed extensive branching morphology, whereas this branching pattern was markedly reduced in RSA59-infected sections (Insets, white arrow). Conversely, A2B5+ OPCs in RSA59-infected sections (Insets) exhibited a more elaborate branching architecture compared with the predominantly perinuclear punctuate Cx47 expression observed in mock-infected sections (Insets).

### 5. RSA59 Infection Downregulates Oligodendrocytic Cx47 Expression in Primary OPCs and Mature OLGs (In Vitro)

The *in vivo* observations were further extended using a reductionist *in vitro* approach to investigate the consequence of RSA59 infection on the oligodendrocytic GJ protein Cx47 in isolated primary OPCs and mature OLGs.

To understand whether alteration of Cx47 *in vivo* could be reflected *in vitro*, primary OPCs and mature OLGs were infected with RSA59 at an MOI of 1. At 24 h p.i., cells were fixed and processed for immunofluorescence analysis. For double labelling, cells were stained with lineage-specific markers (A2B5 for OPCs and CC1 for mature OLGs; Fig. 7, green) along with Cx47 (Fig. 7, red). No significant changes in overall Cx47 distribution pattern were observed in RSA59-infected OPCs (Fig. 7B, Insets) compared with mock controls (Fig. 7A, Insets). In contrast, a marked reduction in branched architecture was observed in infected mature OLGs (Fig. 7D, Insets) compared with mock-infected mature OLGs (Fig. 7C, Insets). Quantitative analysis revealed a significant downregulation of Cx47 expression, measured as mean fluorescence intensity (MFI), in both infected OPCs and mature OLGs (Fig. 7E, F). Notably, reduced Cx47 expression was also observed in neighboring RSA59-EGFP-negative cells (both OPCs and OLGs) that were not actively infected (Fig. S2, A & B), suggesting that viral infection may indirectly influence Cx47 expression, potentially through secretion of soluble factors.

**Figure 7.**
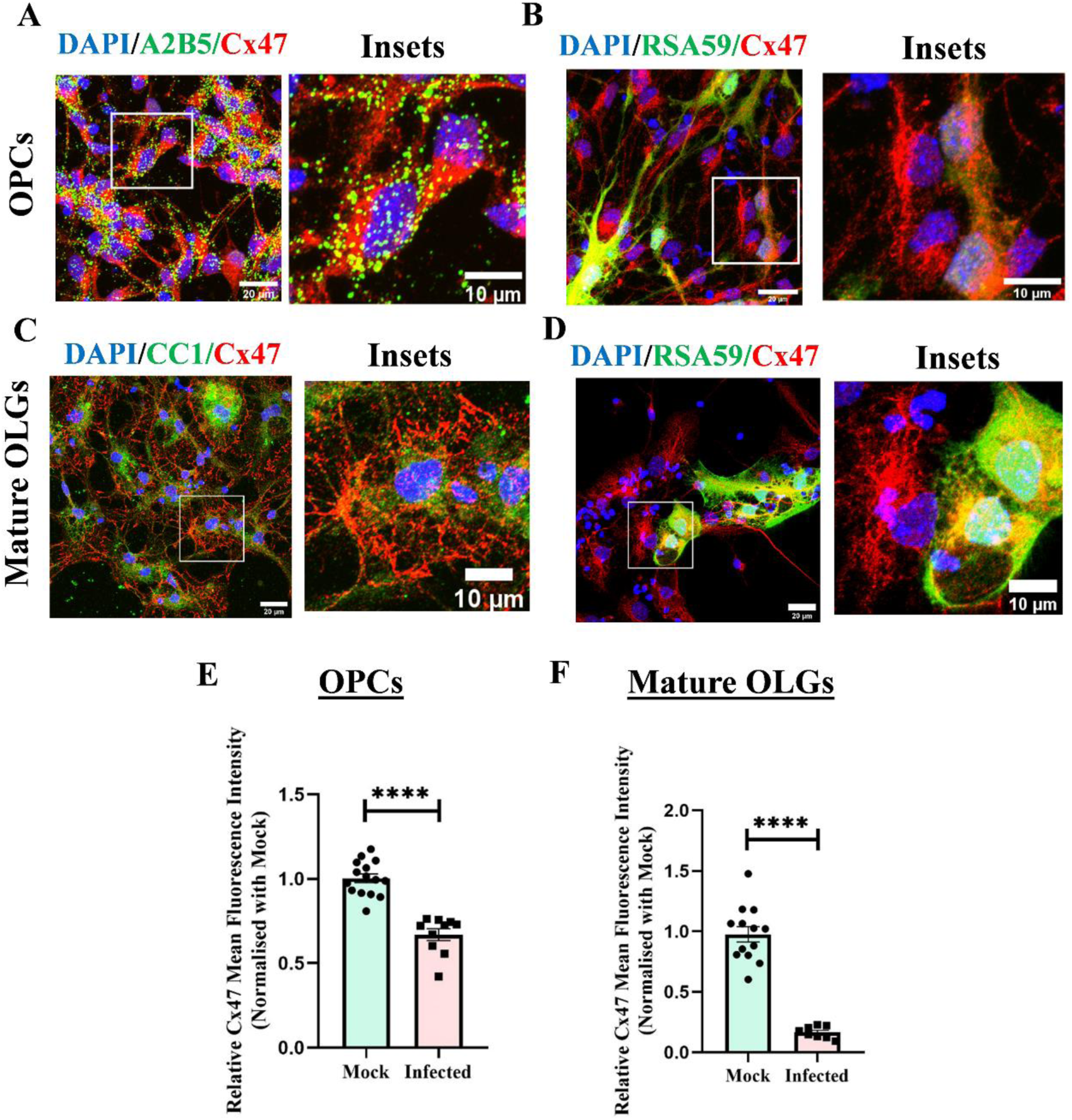
*In vitro* infection of RSA59 downregulates Cx47 expression in primary OPCs and mature oligodendrocytes. (A) Representative confocal images showing Cx47 (red) expression in A2B5 (green) marked OPCs in mock-infected and (B) in 24 h post RSA59 infected (green) OPCs. Insets represent a manually zoomed image of the boxed region in the merged images. Similarly, Cx47 expression in mature oligodendrocytes marked by CC1 (green) and Cx47 (red) in (C) mock-infected mature OLGs and (D) 24 h post RSA59 infected (green) mature OLGs. (E) Quantification of MFI of Cx47 in OPCs and (F) mature OLGs. Calculated in A2B5- and CC1-marked cells for mock- and RSA59-infected (EGFP-marked) cells to specifically check the expression in infected cells. Results were expressed as mean ± SEM (n=3 per group). Asterisks represent statistical significance calculated using unpaired Student’s t-test and Welch correction. P < 0.05 was considered significant. ns - not significant, *P < 0.05, **P < 0.01, ***P < 0.001, ****P < 0.0001

In mature OLGs, the *in vitro* findings were consistent with the *in vivo* results, as a significant decline in Cx47 expression (MFI/cell/field) following RSA59 infection was observed in both systems. In contrast, OPCs displayed differential regulation: decrease in Cx47 expression was observed *in vitro*, whereas a marked increase was detected *in vivo*. Collectively, these findings indicate that oligodendrocytic gap junction protein Cx47 is differentially regulated following RSA59 infection, depending on oligodendroglial lineage (OPCs versus mature OLGs) and experimental context (*in vitro* versus *in vivo*).

### 6. Altered regulation of LIF-LIFR/gp130 signaling axis in the brain upon RSA59 infection

Given that RSA59 infection induces apoptotic cell death of mature OLGs and that LIF-LIFR/gp130 signaling is an established oligodendrocyte survival pathway during immune-mediated demyelination, we next investigated RSA59 also modulates LIF-LIFR/gp130 signaling axis in the brain.

Our result demonstrated that virus (RSA59) infection induces LIF mRNA expression in the brain during the acute stage (d5 p.i.), which gradually decreases towards acute chronic (d10) and chronic phase (d30 p.i.) (Fig. 8A). In parallel, IL-6 an antagonistic cytokine of LIF (also uses gp130 as a co receptor for downstream signaling) showed elevated expression during all the stages post RSA59 infection (Fig. 8B). Furthermore, to determine the source of LIF among the neuroglial cells in the CNS, we employed a reductionist primary cell culture approach. An elevated level of LIF mRNA and protein expression was observed in primary astrocytes upon RSA59 infection for 24 hr as compared to mock-infected control (Fig 8C & D). In addition, we also validated that astrocytes are the major source of mouse LIF (mLIF) production (98-120 Pg/mL) upon RSA59 infection, unlike oligodendrocytes and neurons, where the mLIF levels remain undetectable (Fig. 8E).

**Fig. 8.**
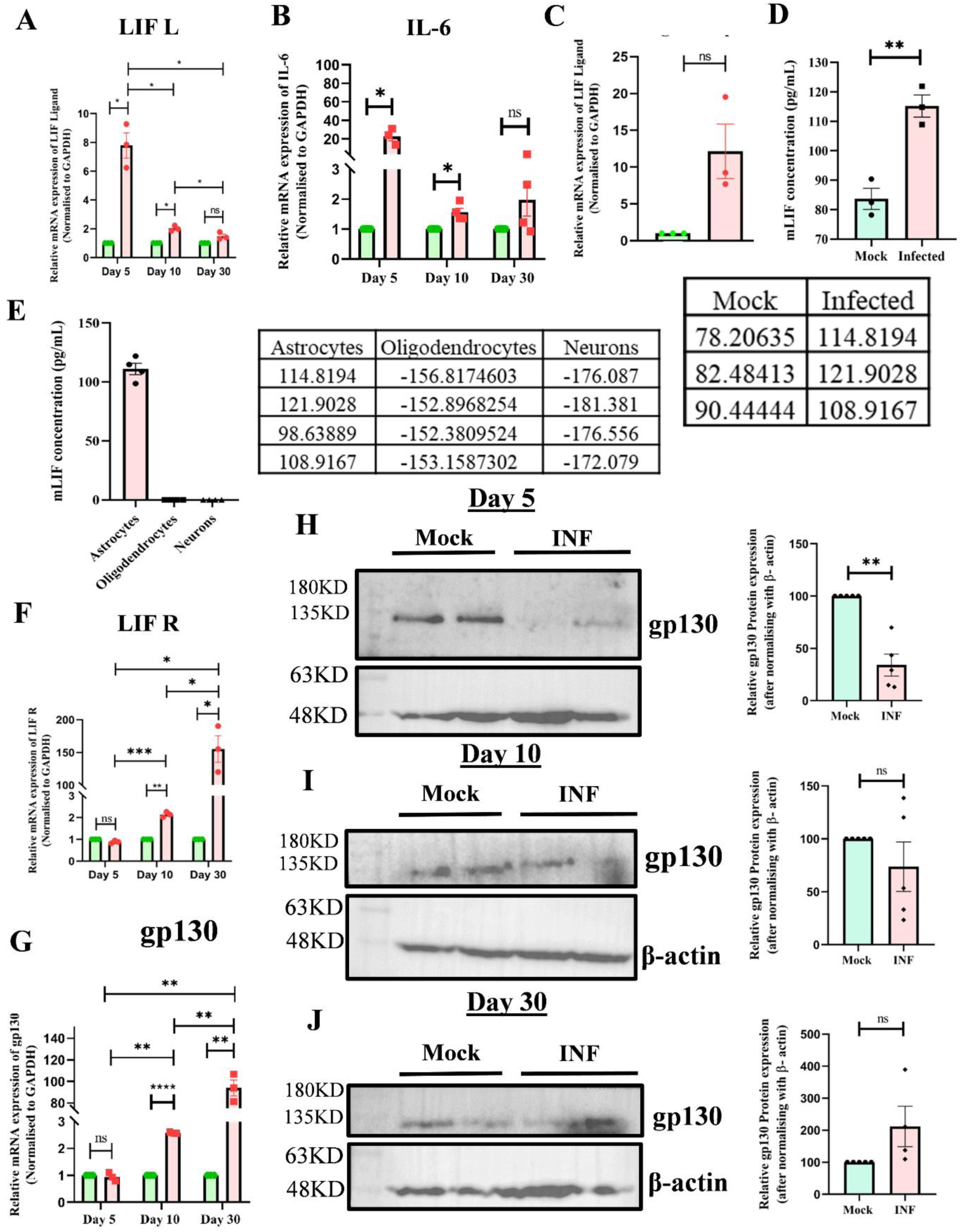
Murine Coronavirus (RSA59) infection differentially modulates LIF-LIFR/gp130 signaling axis in the brain. (A & B) Scatter plots depicting the mRNA expression of LIF and IL-6 in the brain during d5, d10 and d30 post RSA59 infection. (C) The graph indicates mRNA expression of LIF in primary astrocytes 24 hr post RSA59 infection. (D) The Scatter plot depicts the mouse LIF (mLIF) protein concentrations (pg/mL) in cell culture supernatant derived from both mock-infected and RSA59 infected murine primary astrocytes at 24 hr p.i. (E) Similarly, a comparative graph showing mouse LIF (mLIF) protein concentration in the culture supernatant derived from astrocytes, oligodendrocytes and neurons post 24 hr RSA59 infection. Absolute concentrations (in pg/mL) are given in the table. (F & G) Quantification graph showing mRNA expression of LIF-R and gp130 during all the stages (d5, d10 and d30) post RSA59 infection. (H-J) Representative Immunoblot and corresponding scatterplots showing gp130 expression in the brain during acute, acute-chronic and chronic stages post RSA59 infection. Results were expressed as mean ± SEM (n=3-6 per group). Asterisks represent statistical significance calculated using unpaired Student’s t-test and Welch correction. P < 0.05 was considered significant. ns - not significant, *P < 0.05, **P < 0.01, ***P < 0.001, ****P < 0.0001

In contrast to LIF Ligand expression, LIF receptors (LIF R and gp130) mRNA expression was not significantly changed during the acute (d5) phase of RSA59 infection. However, the receptor expression gradually increased towards d 10 p.i. and d 30 p.i. (Fig. 8F & G). Immunoblot analysis of gp130 protein expression showed significant downregulation during the acute (d5) phase of RSA59 infection, but it gradually increases towards d10 and d30 p.i. similar to the mRNA studies. (Fig. 8 H-J). Collectively, our results indicate that RSA59 infection differentially modulates LIF and LIF R/gp130 expressions in the brain.

The observed differential expression of LIF L and LIF R/gp130 post RSA59 infection indicates virus (RSA59) infection may modulate this crucial oligodendrocyte survival signaling pathway, potentially contributing to virus-induced oligodendroglial death and damage.

### 7. Murine Coronavirus (RSA59) Infection Differentially Modulates the ERK Signaling Pathway in a Stage-Dependent Manner

ERK is a key downstream effector molecule of the LIF-LIFR/gp130 signaling, also critical for oligodendrocytes’ survival, proliferation, and maturation (21, 41, 42). Therefore, subsequently we evaluated the effect of RSA59 infection on ERK signaling in the brain.

To investigate the impact of RSA59 infection on ERK signaling pathways, Phosphorylated ERK (pERK) protein expression (indicates activation of ERK signaling pathway) was assessed by immunoblotting using total protein extracts isolated from mock-infected and RSA59-infected brain tissues collected at day 5, day 10, and day 30 p.i. Equal amounts of total protein were resolved and probed with an anti-pERK antibody (Table 1), followed by probing with an anti-total ERK antibody (Table 1) as an internal control. Immunoreactive bands were detected using HRP-conjugated secondary antibodies. The results demonstrated that pERK levels were significantly upregulated in RSA59-infected brain samples during the acute–chronic transition phase (day 10 p.i.) compared with mock-infected controls (Fig. 9C and 9D). In contrast, no significant alterations in pERK expression were observed during the acute phase (day 5 p.i.; Fig. 9A and 9B), and a significant downregulation of pERK expression was observed (Fig. 9E and 9F) during the chronic phase (day 30 p.i.) following RSA59 infection. Overall, the results indicate that RSA59 infection in the brain modulates ERK signaling in a stage-dependent manner. Given the involvement of ERK signaling in regulating various aspects of oligodendroglial function, this differential modulation of the ERK signaling pathway upon RSA59 infection may have diverse functional consequences for oligodendroglial cells.

**Fig. 9.**
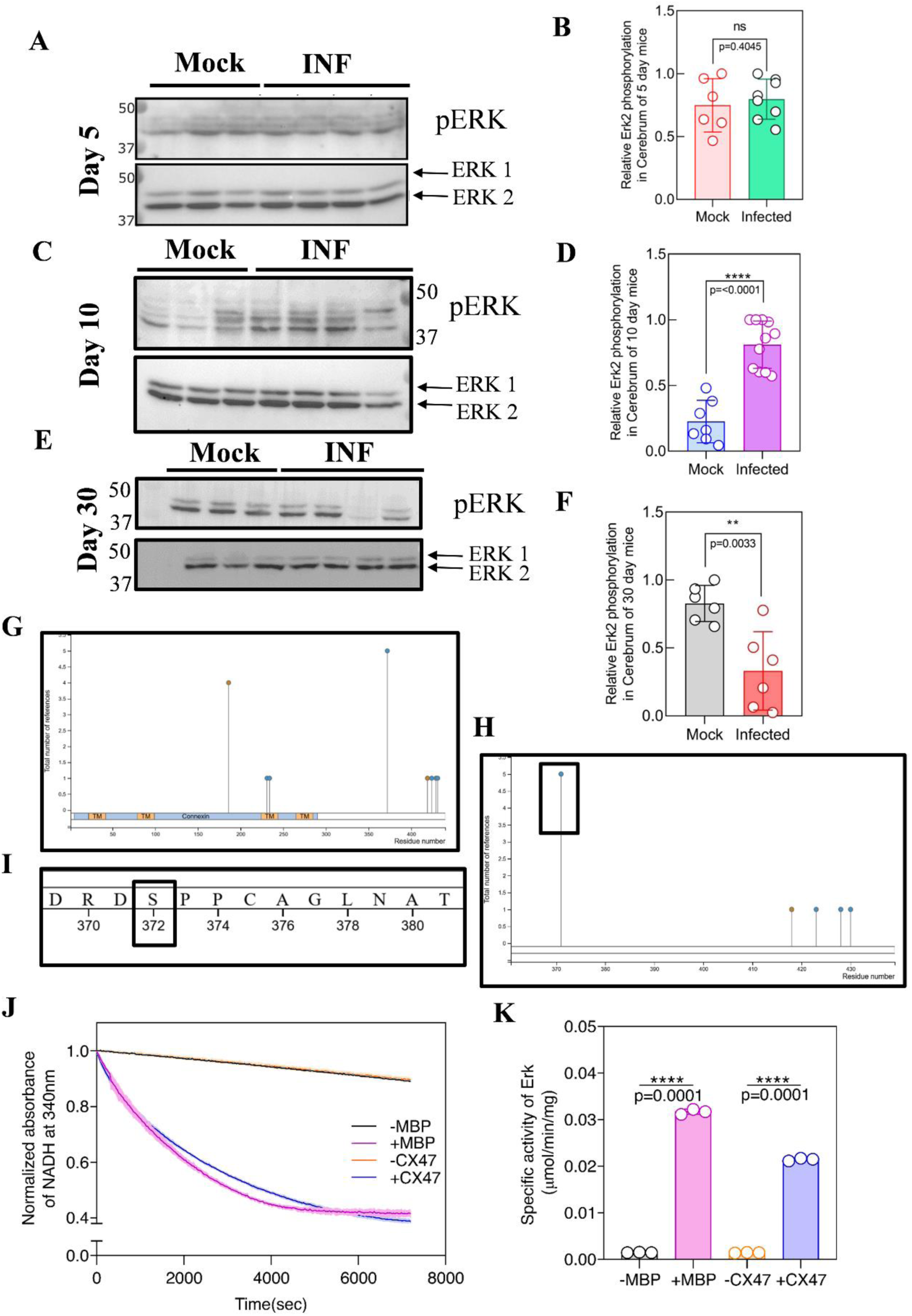
ERK signaling is differentially modulated by RSA59 infection in the brain and identification of an ERK phosphorylation site at Serine-372 position of the Cx47 C-terminal domain. (A, C, E) Representative munoblots and (B, D & F) corresponding scatter plots depicting protein levels of pERK at d5, d10, and d30 p.i. of brain tissues of RSA59 infected mice compared with age matched mock-infected control group. Note that significant upregulation of pERK protein expression during acute-chronic transition phase (d10 p.i), however, no significant changes were observed during d5 p.i. and a significant downregulation was observed at d30p.i. ERK1/2 was used as a loading control. The results were expressed as mean ± SEM (n = 6-8 animals/per group). (G) Four phosphorylation sites at Serine-372, 424, 431 and Threonine-429 of Murine Cx47 (GJC2)-C terminus as indicated in the bioinformatic database PhosphoSitePlus. (H & I) Serine-372 position at Cx47 C terminus is the most reported phosphorylation site. (J) In vitro kinase assays were carried out using activated ERK2 in the presence of either Myelin Basic Protein (MBP), a traditional ERK2 substrate, or a CX47 C-terminal peptide. Phosphorylation reactions were measured using an NADH-coupled assay, where ATP consumption in substrate phosphorylation is coupled to NADH oxidation. The resulting decrease in NADH absorbance at 340 nm was measured over time to determine the rate of phosphorylation. (K) Initial velocities were determined from the slope of absorbance change and used to calculate specific activity (µmol/min/mg) for each substrate. MBP was used as a positive control to confirm ERK2 activity, and the ability to phosphorylate the CX47 peptide confirms its suitability as an ERK2 substrate. Data are mean ± SEM of three independent experiments. Asterisks represent statistical significance calculated using unpaired Student’s t-test and Welch correction. P < 0.05 was considered significant. ns - not significant, *P < 0.05, **P < 0.01, ***P < 0.001, ****P < 0.0001

### 8. Extracellular Signal-Regulated Kinase 2 (ERK2) Phosphorylates at a Specific Serine Residue (Serine-372) of the Cx47 C-terminal domain (Cx47CT)

As demonstrated earlier, RSA59 infection differentially regulated Cx47 expression both *in vivo* and *in vitro* (Result Section 4). Cx47, Non-phosphorylated isoform (I2), and total mRNA levels were increased, whereas phosphorylated isoform (I1) was persistently reduced, suggesting that RSA59 infection may affect Cx47 phosphorylation. Furthermore, we observed that RSA59 infection differentially regulates ERK signaling in the brain, raising the possibility of a functional kinase-substrate relationship between ERK and Cx47. This hypothesis was further supported by a recent report by Latchford L. P. et al. (2025) (34), which demonstrated that all three MAPKs {extracellular signal-regulated kinase (ERK), p38, and c-Jun N-terminal kinase (JNK)} exhibit substrate specificity toward the Cx43 C-terminus, the astrocytic GJ coupling partner of Cx47. Collectively, these findings led us to investigate a potential association of Cx47 with ERK.

Therefore, Cx47 phosphorylation status was further evaluated using a bioinformatic database, PhosphoSitePlus (43). Murine Cx47 (GJC2) contains four documented phosphorylation sites: Serine-372, 424, 431, and Threonine-429 (Fig 9G). Among these, Ser372 is the most frequently reported phosphorylation site (Fig 9H). Although multiple phosphorylation sites have been reported, the specific kinases responsible for phosphorylating Cx47 have not been identified.

To further explore this, a mimetic peptide based on the C-terminal domain of CX47 (DRDSPPCAGLNAL), including the Ser372 residue (Fig. 9I) was employed, and the substrate specificity of ERK2 was examined in a coupled kinase assay. Autophosphorylated ERK2 was produced by co-purification with a constitutively active MEK1 mutant. Using myelin basic protein (MBP) as a standard substrate, the specific activity of ERK2 was found to be 0.03 ± 0.00045 µmol/min/mg. It was then assessed whether the C-terminal peptide of CX47 could act as a substrate for ERK2. The specific activity was found to be 0.02 ± 0.00025 µmol/min/mg (Fig. X). These results show that the C-terminal domain of CX47 is efficiently phosphorylated by ERK2 and thus provide evidence for CX47 as a new physiological substrate of ERK2.

## Discussion

The present study provides a comprehensive mechanistic dissection of the molecular consequences of murine β-coronavirus RSA59 infection on oligodendrocyte biology in the brain. The study specifically focuses on gap junction protein Cx47 dynamics, oligodendrocyte apoptosis, and the LIF-LIFR/gp130-ERK signaling axis. The key findings of this study are: (i) RSA59 directly infects both OPCs and mature OLGs in vitro and in vivo; (ii) RSA59 infection induces mature OLG apoptosis predominantly during the acute phase; (iii) RSA59 infection differentially regulates Cx47 expression in a cell type-specific and disease stage-dependent manner, with persistent increased expression of the non-phosphorylated isoform (I2) accompanied by decreased expression of the phosphorylated isoform (I1); (iv) the LIF-LIFR/gp130 signaling is temporarily dysregulated during RSA59 infection, with early LIF induction by reactive astrocytes that is uncoupled from the receptor downregulation during the acute phase; (v) ERK signalling is differentially modulated across disease stages; and (vi) ERK2 directly phosphorylates the Cx47 C-terminal domain at Ser-372 residue, establishing a novel kinase-substrate relationship that mechanistically links ERK signaling dysregulation to alterations of Cx47 phosphorylation during viral infection.

Our observation that RSA59 directly targets OPCs and mature OLGs aligns with our previous reports (37) and findings from other studies, supporting the notion that direct oligodendroglial infection is a defining feature of demyelinating strains (MHV-A59) and a prerequisite for white matter pathology (44). In contrast, the non-demyelinating strain (RSMHV2) fails to infect differentiated mature OLGs and its white matter spread is restricted (37). Building upon this foundation, the present study demonstrates that RSA59 tropism for oligodendrocytes is also accompanied by robust mature OLG apoptosis in the brain, as evidenced by CC1/TUNEL co-positivity across all disease stages. Murine coronavirus-induced oligodendrocyte apoptosis proceeds via caspase-dependent mechanisms, including Fas/FasL activation (45) and the mitochondrial pathway (46). The predominance of apoptosis at the acute neuroinflammatory phase (day 5 p.i.) observed in the current study implies that direct viral cytopathic effects, rather than solely immune-mediated mechanisms (47), drive the initial wave of oligodendroglial death. The subsequent progressive demyelination at day 30 p.i., despite declining productive viral infection but persistent viral mRNA (48–51), is consistent with a model in which early apoptotic depletion of mature OLGs (52), combined with impaired remyelination (53) associated with chronic Cx47 dysregulation in oligodendrocytes, collectively contributes to chronic demyelination pathology (10, 28, 31).

The sustained downregulation of mature and myelinating OLGs markers (CNPase and MBP) alongside the persistent upregulation of the OPC marker A2B5 across all disease stages is a compelling finding of this study. The increase in A2B5+ OPCs likely reflects an endogenous reparative response, analogous to the OPC proliferative response documented in other demyelinating diseases (54). However, the failure of this expanded OPC pool to effectively remyelinate, as evidenced by persistent loss of myelin markers, mirrors observations in chronic MS lesions, where OPC recruitment is preserved but their differentiation into mature, myelinating OLGs is impaired (55). In this context, the chronic Cx47 dysregulation observed in the current study, specifically the persistent downregulation of Cx47 expression in mature OLGs (PLP+/CC1+) may mechanistically contribute to impaired maturation, as intercellular communication through Cx43-Cx47 heterotypic GJs is essential for the metabolic support and maturation of newly forming oligodendrocytes (56, 57).

The differential regulation of two Cx47 isoforms upon RSA59 infection, with persistent reduction of the putative phosphorylated isoform (I1) and upregulation of the non-phosphorylated isoform (I2), is a novel and important finding of the current study.

Phosphorylation is a well-established post-translational mechanism regulating connexin trafficking, GJ assembly, channel gating, and stability at the plasma membrane (33). Cx43 phosphorylation by various kinases, including MAPK/ERK, controls its redistribution, internalization, and degradation during cell stress and viral infection (58, 59). However, the relevant phosphorylation sites of Cx47 and responsible kinases have not been identified. The PhosphoSitePlus database (43) identifies Serine-372 as the most frequently reported phosphorylation site on Cx47; however, the upstream kinase responsible for this modification remains unknown. Our coupled kinase assay data now provide direct experimental evidence that ERK2 phosphorylates the Cx47 C-terminal peptide encompassing Ser-372. The study identifies Cx47 as a physiological substrate of ERK2 for the first time, representing a previously unreported finding. These results suggest that the stage-dependent ERK alterations observed during RSA59 infection, specifically the downregulation of pERK at day 30p.i. may directly contribute to the persistent loss of Cx47 phosphorylation (I1) and thereby disrupt Cx47 stability, GJ plaque assembly, and intercellular communication in mature OLGs.

The LIF-LIFR/gp130 signaling studies presented here reveal a temporal uncoupling between LIF ligand production and receptor expression during RSA59 infection that has important implications for oligodendrocyte survival. LIF is a major oligodendrocyte survival factor in the context of immune-mediated demyelination; its endogenous production limits mature OLG death during both EAE and spinal cord injury, and exogenous LIF administration promotes OPC proliferation and enhances remyelination (20, 21, 42). Our data establish astrocytes as the primary cellular source of LIF upon RSA59 infection, with OLG and neurons producing no detectable mLIF. This is consistent with the observed reactive astrocyte phenotypes that develop during MCoV infection (60), as well as with reports that reactive astrocytes increase LIF expression in response to CNS injury (61, 62). The acute upregulation of LIF at day 5 p.i. specifically when mature OLG apoptosis is maximal, may represent an inadequate protective response, compounded by the simultaneous downregulation of gp130 receptor expression at this stage, which may reduce the capacity of oligodendrocytes to respond to the LIF-mediated survival signal. The subsequent recovery of LIFR and gp130 expression at day 10 and 30 p.i. may restore LIF-dependent survival signaling at later stages, but by this time the primary apoptotic death has already depleted a significant proportion of the mature OLG populations.

The observed stage-dependent biphasic ERK modulation, specifically, significant upregulation at day 10 p.i. and downregulation at day 30 p.i likely reflects a complex interplay between active immune-mediated demyelination and subsequent failure of remyelination. ERK activation at day 10 p.i. may represent a compensatory response to the LIFR/gp130 recovery occurring at this stage, consistent with the known role of ERK as a downstream mediator of gp130 signaling (63–65). However, while ERK activation can promote OPC proliferation, its sustained activation may paradoxically inhibit OPC differentiation into mature myelinating OLGs, as inhibition of MEK/ERK signaling has been shown to robustly enhance OPC to OLG differentiation and to promote remyelination in EAE and cuprizone models (63, 64, 66). The subsequent chronic suppression of ERK at day 30 p.i. could impair OLG survival signaling, including LIF-mediated neuroprotection, and thereby inducing mature OLG loss and chronic demyelination. The identification of Cx47 as a direct ERK2 substrate provides a molecular explanation for how this dynamic ERK dysregulation translates into altered Cx47 phosphorylation and impaired gap junction function.

These findings have several significant implications. First, we identified the ERK2-Cx47 kinase-substrate axis as a novel mechanistic link between MAPK signaling and oligodendrocytic GJ regulation, which may be relevant not only in viral demyelinating disease but also in MS and other acquired demyelinating conditions where both ERK signaling (67) and Cx47 expression are disrupted (10, 68). Second, the temporal uncoupling of LIF-LIFR/gp130 signaling during the acute phase of infection, when oligodendrocyte apoptosis is maximal, suggests that therapeutic supplementation with exogenous LIF or gp130 agonists during the early stages of viral CNS infection may be particularly beneficial in limiting oligodendrocyte death and subsequent demyelination. Third, the identification of astrocytes as the primary source of LIF production during RSA59 infection, together with prior studies showing that pharmacological restoration of astrocytic Cx43 and ER chaperone ERp29 function with 4-PBA reduces viral spread and demyelination (30), positions astrocytes as central orchestrators of both the pathological (via viral spread and Cx43 dysfunction) and the potentially protective (via LIF production) responses to MCoV infection in the CNS.

Taken together, the current study provides the first comprehensive evidence that murine β-coronavirus RSA59 infection in the brain induces disruption of multiple oligodendrocyte functions, encompassing direct oligodendroglial apoptosis, differential expression of Cx47 gap junction protein at the levels of isoform-specific phosphorylation and cell-type-specific expression, dysregulation of the LIF-LIFR/gp130 oligodendrocyte survival signaling axis, and stage-dependent modulation of ERK signaling. Mechanistically, our study identifies Cx47 as a direct substrate of ERK2 via phosphorylation of the C-terminal Ser-372 residue, providing a novel molecular mechanism linking virus-induced ERK dysregulation to Cx47 post-translational modification and impaired oligodendrocytic gap junction function. These findings substantially advance the mechanistic understanding of virus-induced demyelination and provide a framework for identifying novel therapeutic targets, including the LIF-LIFR/gp130 axis, ERK2-Cx47 phosphorylation, and connexin-mediated glial coupling for the treatment of inflammatory demyelinating diseases such as multiple sclerosis.

## Acknowledgement

We thank the Council of Scientific and Industrial Research (CSIR) and DST for providing fellowships to SKS and RJ, respectively, and IISER Kolkata for providing fellowship support to SG and institutional assistance to Anshita and Ankita during their BSMS studies. We acknowledge the support of the Department of Biological Sciences, IISER Kolkata, the Central Imaging Facility, and Mr. Ritabrata Ghosh. We thank the Small Animal Facility (SAF), IISER Kolkata, for providing the animals and experimental facilities. We sincerely thank Prof. Arnab Gupta, IISER Kolkata, for the Wellcome Trust-Leica SP8 confocal platform and Mr. Brindaban Majhi for his help during image acquisition. We are grateful to Prof. Rahul Das and his laboratory for providing essential experimental reagents. We also thank Prof. Judith B. Grinspan, Children’s Hospital of Philadelphia (CHOP), Philadelphia, PA, for providing MBP, PLP, Gal C, A2B5, and CNPase antibodies.

## Author Contributions

Conceptualization: JDS and SKS, Methodology: SKS, RJ, SG, AM, AM and JDS, Investigation: SKS, RJ, SG, AM, and JDS, Writing: SKS and JDS, Writing-Review and Editing: SKS and JDS, Funding acquisition: JDS, Resources: JDS, Supervision: JDS.

## Ethics approval

All animal experiments conducted in this study were approved by the Institutional Animal Ethics Committee (IAEC) of the Indian Institute of Science Education and Research (IISER) Kolkata and were carried out in strict accordance with the prescribed institutional guidelines. The approved protocols for primary cell culture were IISERK/IAEC/AP/2019/29.01 and IISERK/IAEC/AP/2019/29.02, while the protocols for in vivo experiments were IISERK/IAEC/AP/2024/125 and IISERK/IAEC/AP/2026/182. All experimental procedures were performed in accordance with the statutory rules and regulations laid down by the Committee for the Control and Supervision of Experiments on Animals (CCSEA), Government of India.

## Data Availability

All data supporting the conclusions of this study are presented within the manuscript and its supplementary materials. The data related to the study findings can be obtained from the corresponding author upon reasonable request.

## Supplementary Figures

**Supplementary Figure 1.**
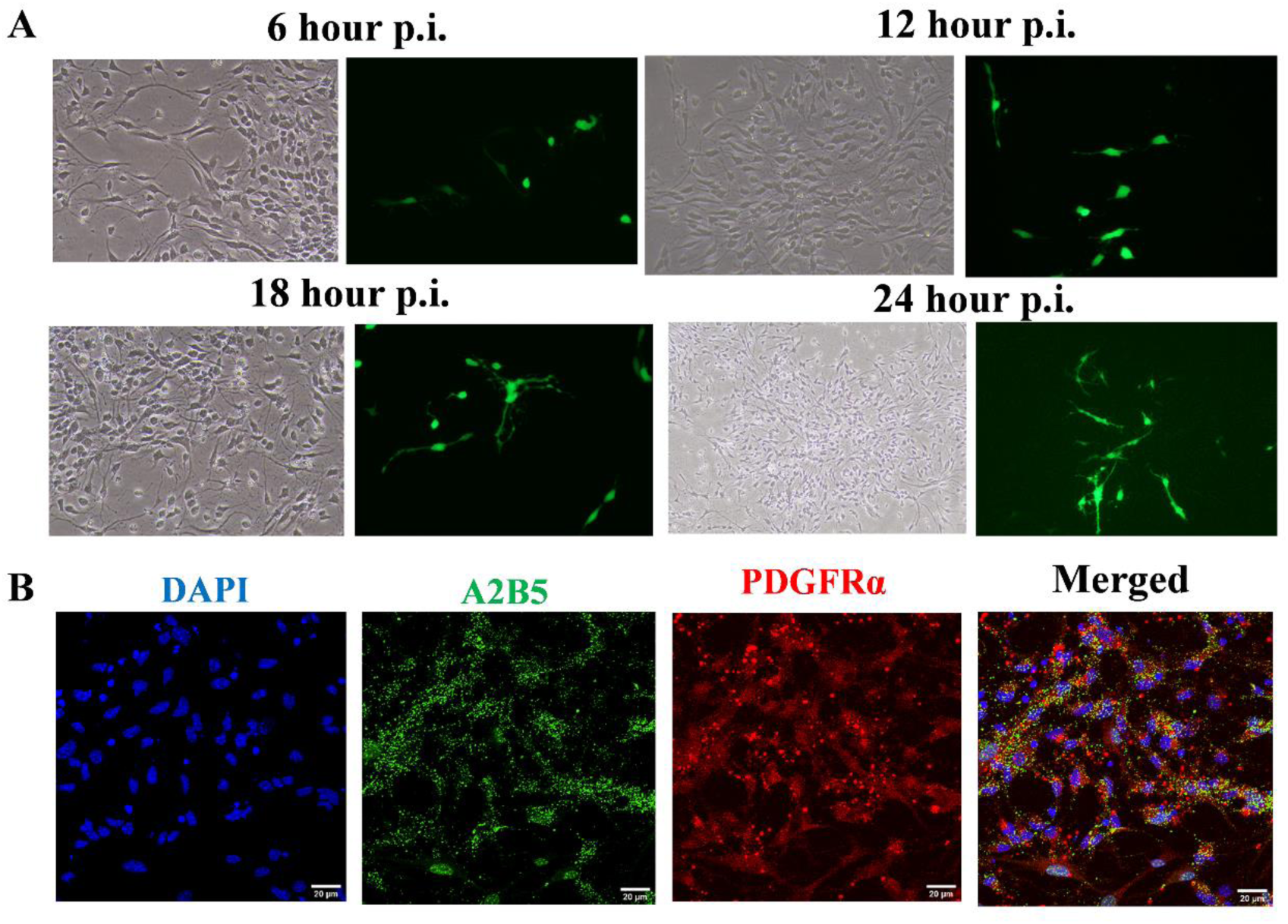
(A) RSA59 infection of primary OPC’s from 6hr p.i. to 24 hr p.i. (B) Characterization of OPC’s using cell specific marker anti-A2B5 and PDGFR α.

**Supplementary Figure 2.**
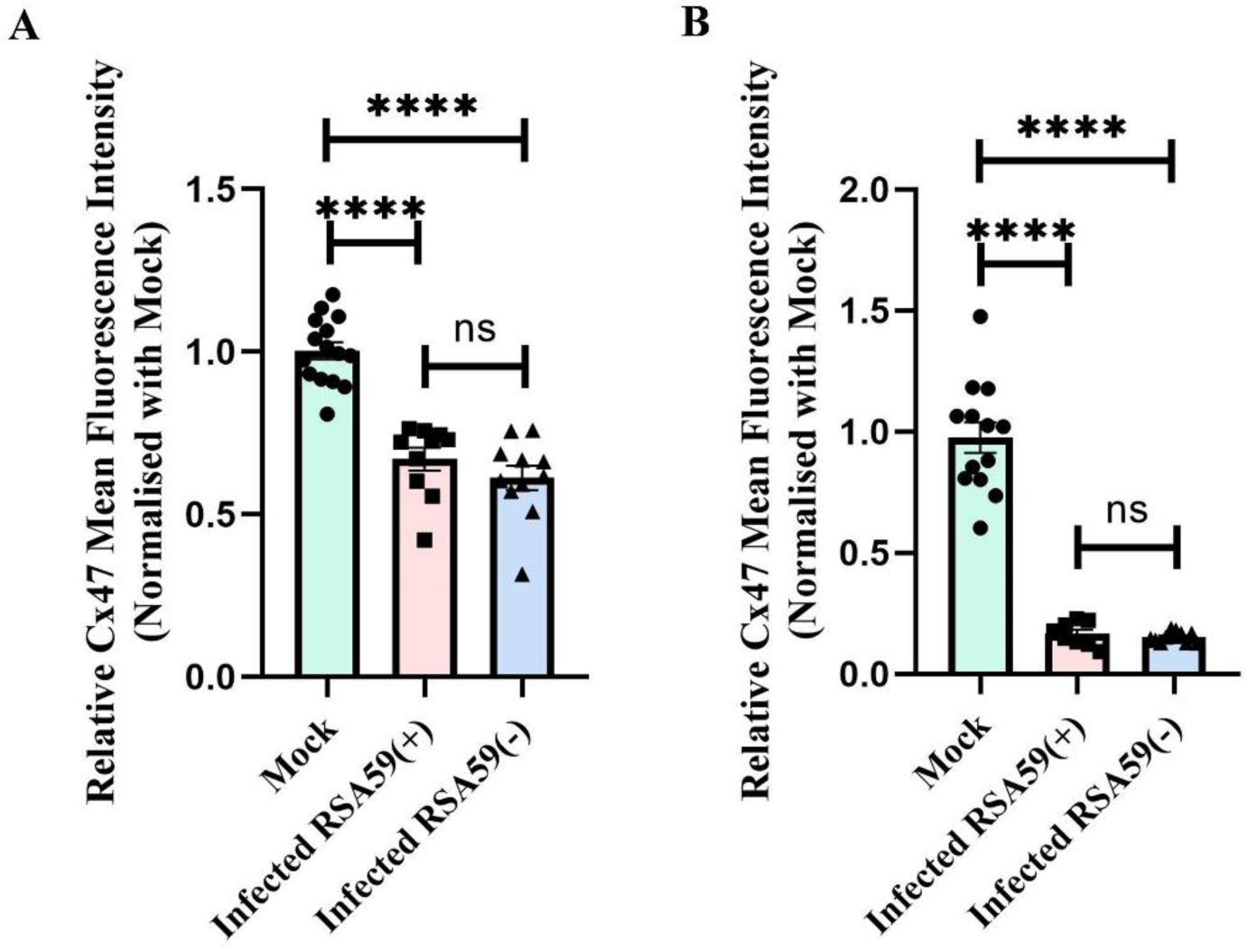
(A) Scatter plot showing Mean Fluorescence Intensity (MFI) of Cx47 in both RSA59 EGFP^+^ OPCs and RSA59 EGFP^−^ OPCs (not actively infected with RSA59 but present in the vicinity) (B) Similarly MFI of Cx47 in both RSA59 EGFP^+^ OLGs and RSA59 EGFP^−^ OLGs

**Supplementary Figure 3.**
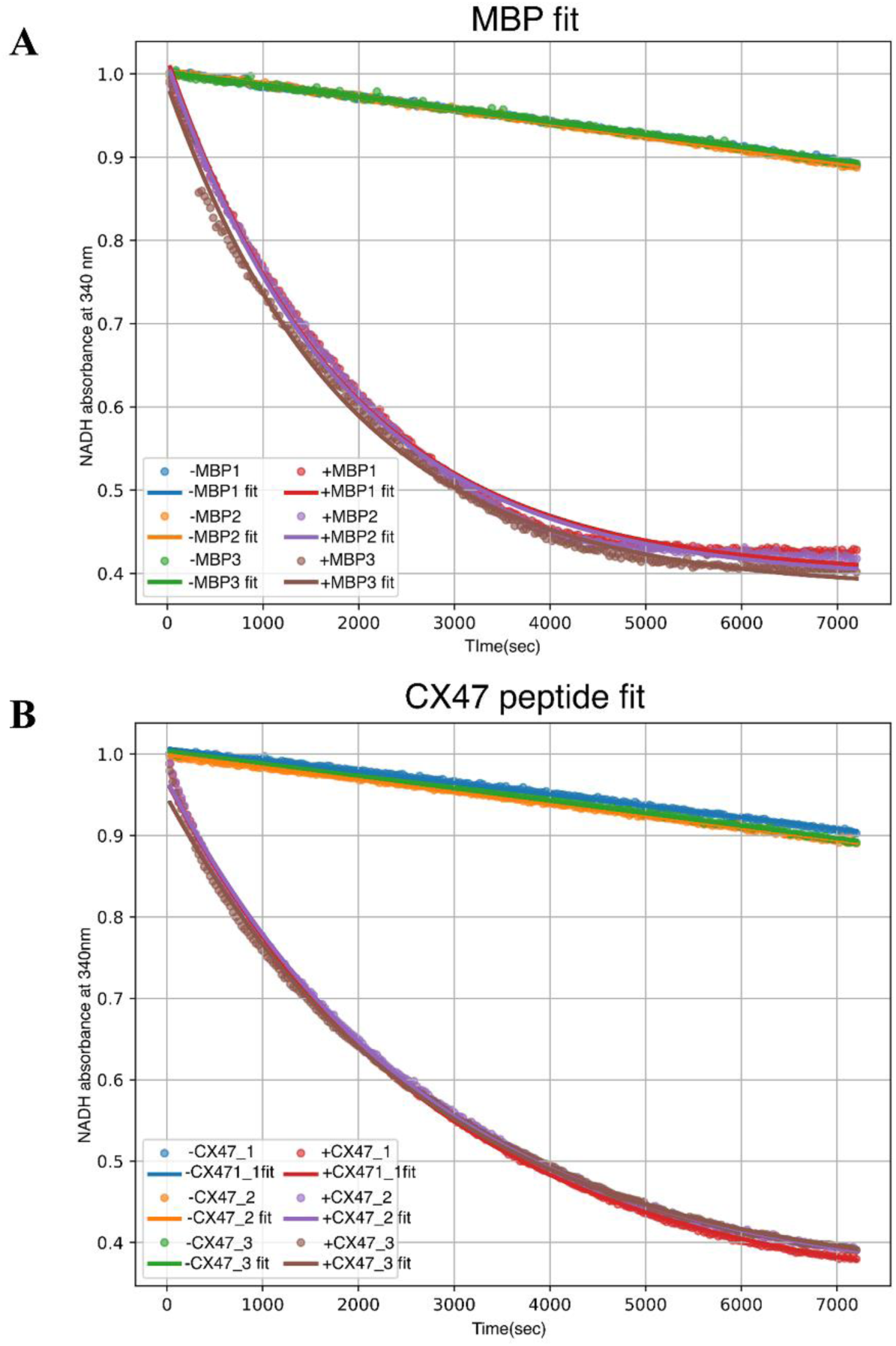
(A) Representative exponential decay fit of NADH absorbance at 340 nm over time for the in vitro kinase assay performed with activated ERK2 using Myelin Basic Protein (MBP) as substrate. (B) Exponential decay fit of NADH absorbance for the assay performed with the CX47 C-terminal peptide as substrate.

